# Population-specific selective sweeps contribute to the maintenance of genetic structure in *Chlamydomonas reinhardtii*

**DOI:** 10.64898/2026.06.19.733452

**Authors:** Jimmy Issa, Scott Ford, Alex N. Nguyen Ba, Rory J. Craig, Rob W. Ness

## Abstract

Microbial eukaryotes often exhibit large effective population sizes and broad dispersal, yet the extent of population structure and the forces shaping it remain poorly understood. While biogeographic structure is often attributed to limits on dispersal, the role of natural selection in maintaining differentiation has received less attention. We investigated population structure and adaptive evolution in the cosmopolitan soil alga *Chlamydomonas reinhardtii*. In addition to the 35 available North American genome sequences we have sequenced 38 new isolates from Ontario (Canada). Population genetic structure analyses demonstrate that these new isolates from Ontario represent a second well-sampled genetically distinct cluster, and that this structure persists despite the presence of recent migrants. Using this data set we were able to conduct genome-wide scans for selective sweeps across the species and within each population. Our results conservatively identify 151 species-wide sweeps and 325 population-specific signals, showing that positive selection is widespread and common. The continued presence of the two distinct genetic clusters as well as loci under differential selection, provide evidence that local adaptation persists despite ongoing gene flow. Together, our results demonstrate that selection plays a central role in reinforcing geographic structure in this highly dispersive microbe.

## 1 Introduction

Compared with their macroscopic counterparts, microbial eukaryotes are thought to have higher effective population sizes (*N_e_*), higher dispersal rates, and lower extinction rates (O’Malley, 2008). Consequently, much debate has focused on the extent to which dispersal and gene flow influence microbial biogeography and population structure (Caron, 2009; O’Malley, 2008). The ubiquity model, envisioned by Baas Becking’s widely-quoted saying “Everything is everywhere, but, the environment selects” (De Wit & Bouvier, 2006), predicts that most microbes have cosmopolitan distributions (within suitable habitats) owing to their high *N_e_*, frequent dispersal, and access to both active and passive dispersal mechanisms that can circumvent the geographic barriers faced by larger organisms (Custer et al., 2022). As a result, high gene flow is expected to reduce extinction and genetic drift, limiting population differentiation (Fenchel & Finlay, 2003; Fenchel & Finlay, 2004; Finlay & Clarke, 1999). However, geographically structured protist populations have been documented across a wide range of systems, including marine phytoplankton and freshwater flagellates (Rengefors et al., 2017; Škaloud et al., 2024), demonstrating that geographic isolation and dispersal barriers can produce significant population differentiation in microorganisms (Casteleyn et al., 2010). In contrast, the endemicity model suggests that some microbes may form endemic populations, with genetic structure arising primarily from barriers to gene flow (Foissner, 1999; Foissner, 2006, 2008).

However, in light of Becking’s original expression, comparatively little attention has been paid to the role of the environment in shaping positive selection and microbial population structure. Few studies have examined geographically explicit patterns of selection and local adaptation in microbial eukaryotes and empirical evidence for local adaptation remains sparse (Kraemer & Boynton, 2017). If environmental heterogeneity favours different traits across geographic regions, positive selection could maintain population differentiation even in the presence of ongoing migration, an outcome consistent with the ubiquity model’s emphasis on dispersal, while being capable of producing structured populations. Distinguishing whether population structure is maintained by isolation or by selection acting against migrants therefore requires explicit tests for geographically variable selection. If local adaptation to varying environments is commonplace in microbial eukaryotes, the role played by the environment to shape selection could drive the evolution of population structure.

Genome-wide scans for selection are a particularly useful approach to identify adaptive traits in microbes as phenotypic differences are often challenging to observe. Such scans can be used to generate unbiased hypotheses for genomic regions, genes, and traits that have been shaped by adaptive processes. An early use of this approach was between Caribbean and Louisiana populations of *Neurospora crassa*, where two genomic islands were identified that appear to play a role in moderating the proliferation of this organism in environments of differing temperatures (Ellison et al., 2011). Selective sweeps are a well-known footprint of positive selection in the genome, where blocks of the genome that are linked with a beneficial allele are driven to high frequency. Selective sweeps result in locally reduced diversity and extended haplotype blocks around the selected locus. When selective pressures differ across the landscape, sweeps can additionally generate strong differentiation among populations at these loci (Pavlidis & Alachiotis, 2017; Stephan, 2019). As a result, selective sweep analysis has become a common part of the population genetics toolkit in describing natural populations (Bourgeois & Warren, 2021) and a variety of methods have been developed to identify sweeps (Abondio et al., 2022; Weigand & Leese, 2018). Sweep analysis has been applied widely including in fungi (Spanner et al., 2021), strawberries (Hu et al., 2022), sheep (Eydivandi et al., 2021), and in humans (Hernandez et al., 2011; Sabeti et al., 2002). Analyses of population structure and selective sweeps in microbial systems have typically focused on pathogenic and parasitic species that threaten human public health and agriculture (Ailloud et al., 2019; Badouin et al., 2017; Duan et al., 2021; Henden et al., 2018; Huang et al., 2023; Kang et al., 2021; Onetto et al., 2022; Spanner et al., 2021). Despite the growing body of literature on selective sweeps in pathogenic and parasitic microbial species, the role of selection in driving population differentiation of free-living, eukaryotic microbes remains poorly explored. This uncertainty, which also limits efforts to address the debate between ubiquity and endemicity models of microbial population structure, is partly due to limited sampling of these microbes from different geographic regions.

The green alga *C. reinhardtii* is a haploid, unicellular soil microbe, and is a model organism for photosynthesis, plant cell biology, and cilia biology (Salomé & Merchant, 2019). *C. reinhardtii* was first sampled in Massachusetts, and today, most natural isolates are from the eastern USA (including Pennsylvania, North Carolina, Minnesota, and Florida) and Quebec (Canada) (Craig et al., 2019; Flowers et al., 2015), although a small number have also been discovered in Japan (Nakada et al., 2010) and South Korea (Kim et al., 2022). Building on the earlier work of Flowers et al. (2015), we previously performed the most detailed analysis of population structure for this species (Craig et al., 2019), examining 35 isolates of *C. reinhardtii* from North America and two from Japan. We identified population structure distinguishing a group of isolates called North American 1 (NA1), encompassing all but one of the Massachusetts and Quebec isolates, from all remaining isolates in the eastern USA, which were called NA2. This geographic structuring mirrors patterns observed in wild yeast (*Saccharomyces paradoxus*) and other taxa, where lineages are thought to have diverged in isolation in distinct glacial refugia and subsequently come into secondary contact (Charron et al., 2014; Craig et al., 2019). Despite the strong geographic structure, we also previously found evidence for admixture in the genomes of most strains, and identified one strain sampled in Quebec that appeared to be a descendant of a recent migration event from an NA2 population (Craig et al., 2019). Thus, consistent to a degree with both the ubiquity and endemicity models, these two genetically differentiated groups persist despite ongoing migration and gene flow, raising questions about whether local adaptation could be maintaining population structure in the face of dispersal. Since this study, we developed a protocol to identify field isolates of *C. reinhardtii* (Ford et al., 2022) and used it to sample 38 new strains in Ontario, representing the first strains of *C. reinhardtii* discovered in this area. Here, we perform whole-genome re-sequencing on these novel strains combined with previously characterized genome sequenes from North America to conduct population structure analyses. We find that along with the Quebec isoaltes, these new new Ontario isolates represent two well-sampled geographic clusters of *C. reinhardtii*, making it possible to conduct both species-wide and population level analyses of positive selection in this model species (Kraemer & Boynton, 2017). Using this approach, we test the hypothesis that natural selection plays a role in maintaining local microbial population structure.

## 2 Materials and Methods

### 2.1 Field Sampling

Soil samples were collected during the summer of 2022 from sites across the Greater Toronto Area and surrounding regions (Figure S1; Table S1-S3). Sampling locations spanned a spectrum of urban to non-urban habitats that included disturbed soil (tire tracks, footpaths, cultivated gardens, farmer fields, river banks) and natural soil (meadows, forest floor), with both wet and dry soil types represented (Table S1). On each sampling day, temperature, humidity and precipitation were recorded (Table S1).

Identification of *C. reinhardtii* in soil samples was conducted using the method we described previously (Ford et al., 2022). This method was used to identify the first seven Ontario field isolates (BMS-1, UTM7-1, CB6-3, CL3-1, JG4-1, W13-1, and W13-2). Briefly, 7-12 mL soil samples were collected in 30 mL centrifuge tubes and combined with an equal volume of Tris-Acetate-Phosphate (TAP) medium. Tubes were incubated at 25 °C under plant growth lights for 3-6 days until algal growth was visible. A 20-50 µL aliquot of supernatant from each soil sample was thin-spread onto a 1.5% TAP agar plate (6 cm diameter) and incubated for an additional 3-7 days under constant light until green algal colonies >1 mm in diameter were observed. Individual colonies were picked; the bulk of each colony was transferred to a 5% Chelex-100 solution for crude DNA extraction as described (Cao et al., 2009), while residual cells on the tip were used to inoculate liquid TAP cultures. Because a single cell can divide vegetatively during growth in the incubator, we can potentially generate clones in the laboratory that do not represent free-living clones. To avoid oversampling of these clones, we limited our screening to 2-4 green colonies per agar plate, prioritizing colonies that exhibited unique morphological features. Clones that are generated through this process were removed after sequencing if their genomes are nearly identical (see below).

Colony PCR was performed using Chelex-100 crude DNA extracts and primer sets with varying specificity for *C. reinhardtii* and its close genetic relatives (Ford et al., 2022). *C. reinhardtii*-specific primers were used to identify *C. reinhardtii*, whereas the broader-range primer sets were used to identify closely related species (Table S4). PCR conditions were as follows: (1) 94 °C for 2 min; (2) 35 cycles of 94 °C for 30 s, 55 °C for 40 s, and 72 °C for 1 min 15 s; (3) 72 °C for 5 min. For positive colonies, we confirmed species identity by amplifying and Sanger sequencing a fragment of the rbcL gene (ribulose 1,5-bisphosphate carboxylase/oxygenase large subunit). *C. reinhardtii* cultures were maintained in the lab on TAP agar slants (Gorman & Levine, 1965). Streak plating was performed to reduce bacterial contamination in cultures that displayed signs of bacterial or fungal contamination.

### 2.2 Genomic DNA extraction, library preparation, and sequencing

Each *C. reinhardtii* culture was grown on TAP agar plates (5 cm) until a thick lawn formed (4–5 days). DNA was extracted from these cells using a phenol chloroform extraction (Craig et al., 2019). DNA was resuspended in ultrapure water and concentration of DNA samples was determined using a Qubit fluorometer.

Dual-indexed genomic DNA libraries were prepared following the protocol described by Santangelo et al. (2022), with minor modifications. Genomic DNA was diluted to 10 ng/*µ*L in a final volume of 25 *µ*L. DNA was sheared to an average fragment size between 500 and 600 bp using a Bioruptor Pico Sonicator (Diagenode Inc., Denville, NJ, USA) for 8 rounds (20 s. each, with 15 s. rest). Small fragments were removed using a SPRI bead solution (Rohland & Reich, 2012): 0.1% (v/v) carboxyl-modified Sera-Mag Speed-beads, 18% (w/v) PEG-8000, 1 M NaCl, 10 mM Tris-HCl (pH 8.0), 1 mM EDTA (pH 8.0), 0.05% Tween 20) at a ratio of 0.8X bead solution:DNA volume. Libraries were generated in four sequential enzymatic reactions: (1) end repair, (2) A-tailing, (3) adaptor ligation, (4) indexing PCR, each followed by a SPRI bead cleanup step to remove residual primers and dNTPs (Santangelo et al., 2022). End repair was performed using T4 DNA polymerase and T4 polynucleotide kinase with ATP and dNTP, resulting in 5*^′^* phosphorylated blunt-ended fragments. 3*^′^* adenosine tailing was completed using Taq polymerase with dATP. iTru adaptor stubs, which were first prepared by annealing iTrusR2-stubRC and iTrusR1-stub adaptors (Glenn et al., 2019), were then ligated to the DNA fragments using T4 DNA ligase. Next, PCR was performed using iTru5 and iTru7 primers (Glenn et al., 2019) to extend the adaptor stubs, and add unique i5 and i7 indexes, as well as the universal p5 and p7 sequences used for Illumina flow cell hybridization. The PCR was performed using Phusion HiFi polymerase (Thermofisher) with the following reaction conditions: 3∼min at 98 *^◦^*C, (30∼s at 98 *^◦^*C, 30∼s at 65 *^◦^*C, 60∼s at 72 *^◦^*C) *×* 15 cycles, 1∼min at 72 *^◦^*C.

All libraries were quantified on a Qubit using the high-sensitivity assay kit (Thermofisher), and run on a 1.5% agarose gel at 100∼V to visualize library size distribution. Prior to sequencing, individual dual-indexed libraries were pooled at equimolar dilutions to ensure multiplexed sequencing would generate roughly equal coverage of each sample. The pooled library was submitted to The Centre for Applied Genomics at SickKids Hospital (Toronto, Ontario), for sequencing on a NovaSeq X platform using 150bp paired-end reads. A mean of 27,745,133 reads were generated for each sample.

### 2.3 Read alignment, mapping, variant calling, and SNP filtering

Sequence reads for each strain were aligned using bwa mem (version 0.7.17-r1188 (Li & Durbin, 2009)) to the *C. reinhardtii* version 6 genome (Craig et al., 2022). Mean coverage for each sample was ∼40.4. Variants were called for each sorted alignment file (BAM) using GATK Haplotypecaller to create GVCFs with the following non-default options, heterozygosity 0.01, indel-heterozygosity 0.001, min-base-quality-score 10, output-mode EMIT_ALL_CONFIDENT_SITES, sample-ploidy 1, standard-min-confidence-threshold-for-calling 1, emit-ref-confidence BP_RESOLUTION (GATK version v 4.2.6.1 (McKenna et al., 2010)). These GVCFs were combined with CombineGVCFs and final variants were called using GenotypeGVCFs using the same ploidy and heterozygosity priors above, along with the option include-non-variant-sites to enable estimation of diversity levels including invariant sites. Following Craig et al. (2019) for SNP filtering, only invariant and biallelic sites were considered for downstream analysis. We applied filters independently to the genotype calls of each isolate, requiring a minimum depth of three mapped reads and a maximum depth not exceeding the mean depth plus four times its square root, to exclude regions with potential copy number variation. Genotypes within 5 bp of an INDEL, those with a genotype quality (GQ) < 20, or with < 90% of informative reads supporting the call were filtered. Furthermore, we excluded the ∼600kb *mt*+ and *mt−* (mating type) loci.

### 2.4 Identifying clonality and mating type

Across 20,000 random SNP subsamples, pairwise genotype comparisons revealed a pronounced bimodal structure in sequence similarity. One cluster of comparisons showed extremely high similarity, with *≥* 19,990 matching SNPs (*≥* 99.9% identity), differing by only a handful of sites. In contrast, the next closest comparisons exhibited a sharp drop in similarity (19,940 matches), corresponding to an approximately fivefold increase in the number of differences. Beyond this point, pairwise differences increased gradually, reaching a maximum of 573 differences across all comparisons (Figure S2). To determine which member of a clonal pair (or three-way pair, at most) should be retained, the sequencing quality for each isolate was identified and compared using the percentage of genomic sites that meet the condition of Depth *≥* 5. The clonal individual with a higher percentage of sites meeting this cutoff was retained. With this approach, a total of eight clonal sets were identified (Table S5), and from them, ten clonal isolates (MW46-07, MW46-13, MW46-14, MW46-15, MW46-18, HH93-02, PW10-05, PW10-02, PW10-04, PW21-02) were excluded. When run on all isolates, this method also identified that less than 6% of sites in the isolate MW46-11 passed this filter, resulting in it being filtered out of downstream sweep analyses as well. Mating type of each isolate was identified using coverage of the *MID* gene (Table S6), which is only present on the *minus* mating type (Ferris & Goodenough, 1997).

### 2.5 Population structure analysis

Craig et al. (2019) found that then-known North American isolates of *C. reinhardtii* could be assigned to one of two genetic lineages that they denoted as NA1 and NA2. To determine whether the new isolates belong to one of these populations or new populations, we repeated our previous analyses, including setting the same VCF filters as before (Craig et al., 2019) along with additional filtering steps as needed depending on the program used, as described below. No missing data were allowed during filtering, resulting in 1.44 million high-quality SNPs. With this data, three methods were used to infer the number of populations across all 73 North American isolates.

First, the Bayesian clustering program STRUCTURE v2.3.4 (Pritchard et al., 2000) was run. Additional filtering prior to the run included thinning variants to one per 20 kb to avoid linkage disequilibrium. This program takes as input a K value denoting the number of clusters, and attempts to assign ancestry proportions of each individual across that number of clusters using Markov chain Monte-Carlo (MCMC) simulations. Simulations were performed for a range of K values, where the optimal K was identified by maximizing the log-likelihood ln Pr(*X|K*) for the observed genotypic data X. A limitation of this approach is that increasing K past the true number of clusters continues to slightly maximize the ln*P r*(*X|K*) quantity due to overfitting. However, Evanno et al. found that the K value that maximizes the second-order rate of change of the ln*P r*(*X|K*) quantity (a value denoted as ΔK) is better at corresponding to the true K (Evanno et al., 2005; however see Gilbert, 2016; Janes et al., 2017 for limitations). Therefore, the STRUCTURE program was run from K=1 to K=10 with 15 replicates per K value, with 1,500,000 iterations of burn-in and 500,000 iterations of MCMC. This process was parallelized using structure-parallel (Koc, 2021). Results from running STRUCTURE replicates were submitted to STRUCTURE HARVESTER (Earl & vonHoldt, 2012) to find Evanno’s ΔK and the distribution of ancestry proportions of several values of K were visualized using STRUCTURE PLOT (Ramasamy et al., 2014). As Evanno’s K is biased towards achieving a K=2 result (Janes et al., 2017), additional and hierarchical methods are also needed to determine population structure.

Second, patterns of haplotype-sharing across the genome of each individual were inferred using the fineSTRUCTURE program. fineSTRUCTURE identifies, for each haplotype in an individual (recipient) genome, the haplotype that it most immediately coalesces among the other individuals (identified as donor). The result is that the number of shared haplotypes each individual has with all other individuals is determined (made possible by *C. reinhardtii*’s haploid genome), and these results are plotted as a coancestry matrix by the program (Lawson et al., 2012).

Third, a PCA-based approach was used utilizing the Bioconductor package SNPRelate (Zheng et al., 2012). In this approach, the principal components (PCs) of the SNP data are found, and the two PCs which explain most of the variance of the data can be used to plot the PCA. Each individual represents a point on the plot, and clustering points correspond to individuals who have a closer genetic distance to each other than to other individuals and these clusters are often used to identify natural populations.

### 2.6 Diversity, divergence, and differentiation statistics

Three statistics were measured to capture diversity within clusters and differentiation between them: pairwise nucleotide diversity (*π*), nucleotide divergence (d*_xy_*), and F*_st_* (Weir & Cockerham, 1984). All statistics were estimated in 30 kb windows using the Pixy program (Korunes & Samuk, 2021). Naturally, *π* was estimated for all clusters, whereas d*_xy_* and F*_st_* were measured between all unique cluster pairs.

To assess whether diversity patterns were shared between populations, we calculated the Pearson correlation coefficient between Ontario and Quebec *π* estimates across matched 10 kb genomic windows. To test whether neutral nucleotide diversity was reduced near centromeres, we compared mean *π* of 4-fold degenerate sites in a 1 Mb window centered on each centromere midpoint to mean *π* in the remaining chromosome arms. Significance was assessed with a chromosome-level permutation test: for each of 10,000 replicates, a 1 Mb window was placed at a random valid location within each chromosome and the arm-centromere difference was recalculated. The one-tailed *p*-value was the proportion of permutations in which the random placement produced a difference equal to or greater than the observed centromere-associated reduction. To test for subtelomeric diversity suppression, we classified windows within 50 kb of chromosome ends as subtelomeric and compared their *π* distributions to those from chromosome arms using two-tailed Wilcoxon rank-sum tests (Figure S4).

### 2.7 Selective sweep analysis

Two selective sweep statistics were used to identify putative windows which have undergone a selective sweep. The first was the *µ* statistic implemented in the RAiSD v2.0 program (Alachiotis & Pavlidis, 2018; Szpiech, 2024). The *µ* statistic is calculated as a composite of three other selective sweep signatures (denoted *µ_var_*, *µ_SFS_*, and *µ_LD_*) to maximize its ability to detect the sweep signal, and was separately run on each cluster in the analysis and the full species (see Results). *µ* scores are centered with a mean of 1, and larger scores indicate a stronger signal of a selective sweep. A susceptibility of the *µ* statistic is that repetitive regions reduce local levels of genetic diversity by reducing local recombination rates, which in turn inflates the *µ_var_* value used to calculate *µ*, and therefore inflates the entire statistic. For this reason, repeats were first identified by passing a custom repeat library to RepeatMasker (A.F.A. Smit, R. Hubley & P. Green RepeatMasker at http://repeatmasker.org) and supplemented with microsatellite and satellite annotations from Tandem Repeats Finder, as per Craig et al. (2022). All genomic windows where repeat densities were *≥* 60% were filtered when running the program. To eliminate false-positives among resulting windows returned by the program, two approaches were considered. The first was to perform background selection simulations, as background selection can reduce genetic diversity and so inflate *µ_var_* for similar reasons to high repeat density. To do this, the best demographic history was first inferred using the fastsimcoal program (Excoffier & Foll, 2011). Parameters included performing (-n) 500,000 coalescent simulations per demographic scenario with (-L) 80 optimization cycles. Furthermore, 100 replicate runs of the program were performed to try to ensure that the global optimum of the combination of parameters was achieved. The following demographic histories were considered: (i) no historical events (ii) bottleneck prior to divergence of populations (iii) population expansion prior to divergence (iv-v) bottleneck or population expansion in one lineage following divergence (vi-vii) bottlenecks or population expansions in both lineages following divergence (viii) bottleneck in one population and population expansion in the other population following divergence. The best demographic history was chosen as the one which minimized the difference between the maximum likelihood value assuming a perfect fit between the observed and expected joint site frequency spectrum (SFS) and the maximum likelihood estimated by the program from the specified parameters. This demographic scenario was then simulated on 17 fragments of 100 kb chromosomes (to reflect the number of chromosomes of *C. reinhardtii*) under the influence of background selection in SFS_CODE (Hernandez, 2008). Population fitness (*γ* = N*_e_*s) values of *γ* = 50, 75, 100, 200, 1000 were tested with 1000 runs of SFS_CODE for each value. The `–`VCF flag was set to output the results in a VCF and -A to do so without also printing the genomic sequence, and these VCFs were run into RAiSD to determine the distribution of *µ* values appearing under background selection in the best demographic scenario. A second, and commonly used approach in the literature for eliminating false-positives is to take all windows whose *µ* statistic is in an upper percentile of all windows (Feng et al., 2023; F. Ji et al., 2021), which are likely to be enriched in true-positive scores.

Additionally, the XP-nSL statistic (Szpiech et al., 2021) as implemented in the selscan v2.0.0 program (Szpiech, 2024) was also used as a between-population statistic to help identify windows under lineage-specific selection. The selscan program as-is does not accept haploid VCFs. A solution to this, however, is to diploidize the *C. reinhardtii* VCF (turning alleles of state 0 to 0/0 and 1 to 1/1) and then run the program in unphased mode (Zachariah Szpiech, personal communication). After running selscan, the resulting XP-nSL statistic values were normalized prior to downstream analysis using the norm package that is packaged with selscan. Unlike some other cross-population statistics such as XP-EHH, XP-nSL does not require a genetic map to be used. An object and reference population need to be defined (specified by the --vcf and --vcf-ref flags respectively). We were primarily concerned with the comparison between Ontario and Quebec samples and opted to exclude clones and putative migrants from each group to avoid spurious signals due to distortions in patterns of linkage or the site frequency spectrum. In the output, positive scores are indicative of a sweep in the object population, whereas negative scores are indicative of a sweep in the reference population. Prior approaches were followed (Santangelo et al., 2025; Szpiech et al., 2021) to identify windows under selection in a manner that balances maximizing the true-positive rate with minimizing the false-positive rate. In 50 kb windows, the mean XP-nSL score and the proportion of XP-nSL scores *≥* 2 were determined. The top 1% of windows from both distributions were extracted, and windows falling in the top 1% of both distributions were putatively identified as having experienced a selective sweep. For these calculations in each window, positive and negative XP-nSL scores were considered separately so as to prevent positive and negative scores (both acting as signals of a selective sweep but in different populations) from averaging each other out. (This means that for negative scores, smaller means of the XP-nSL score and proportions of XP-nSL values *≤* −2 are considered.) Therefore, candidate windows under selection were independently obtained for both populations from positive and negative scores.

Permutation testing was done for further validation of results. Top positive XP-nSL windows (TPW) and top negative XP-nSL windows (TNW) suggest candidate windows for selective sweeps for each of the two clusters that the comparison was run on. There is a degree of overlap expected between the sweeps localized to each population from XP-nSL, and those derived when RAiSD is run on each of those populations respectively. To test for a non-random association between these outlier genomic regions (windows), the regioneR package (Gel et al., 2016) was used. As an additional control, permutation testing can be done across populations where the prediction is that there is no significant overlap. Bonferroni corrections are applied to correct for running four separate tests.

### 2.8 Functional enrichment

Genes from the v6.1 annotation localized within or partially intersecting windows putatively identified as having experienced a selective sweep in the previous step were extracted using pybedtools (Dale et al., 2011), a Python-wrapper for the command-line BEDtools program (Quinlan & Hall, 2010). An extracted list of genes was uploaded to the g:Profiler web tool for gene ontology (GO) enrichment analysis (Raudvere et al., 2019) when setting the significance threshold to Benjamini-Hochberg FDR. g:Profiler offers prepared Gene Matrix Transposed (GMT) files for separate functional enrichment analysis of Biological Process, Cellular Component, and Molecular Function ontologies using its g:OST tool for a variety of organisms, including *C. reinhardtii*. To use gene annotations of the v6.1 release of the of the *C. reinhardtii* genome (Craig et al., 2022), a custom-input set of GMT files were prepared (Raudvere et al., 2019) for enrichment analysis in g:Profiler (preparation details and custom script located at https://github.com/jimmissa/chlamy_sw/). Results were visualized in the EnrichmentMap add-on to Cytoscape v3.9.1 (Merico et al., 2010; Otasek et al., 2019; Shannon et al., 2003).

## 3 Results

### 3.1 Genetic clustering of new isolates

We reassessed the population structure of North American *C. reinhardtii* to determine how newly sampled Ontario isolates relate to previously described clusters (Craig et al., 2019). Under the framework of allopatric divergence in glacial refugia, contact between the NA1 and NA2 lineages is expected near the St. Lawrence River, and Ontario isolates may therefore be expected to be more closely related to NA2 (spanning most of the eastern USA) than to NA1 (Quebec and northeastern USA).

To re-assess population structure, we first conducted PCA analysis. PCA revealed a primary axis of variation (PC1, 23.0%) that clearly separated isolates from Quebec and Massachusetts (MA) from all other North American samples, consistent with strong genetic differentiation of the NA1 lineage (Figure 1). A secondary axis (PC2, 4.7%) distinguishes Ontario isolates from the remaining samples belonging to NA2. Notably, the two isolates from Pennsylvania (CC-2342 and CC-2344) and the previously identified NA2 migrant strain from Quebec (CC-3079) were positioned intermediate to the Ontario isolates and the remaining NA2 isolates (all of which were sampled more than 800 km from Ontario). One potential migrant Ontario isolate, HH93-02, was located at the extreme of PC2 in closest proximity to the more geographically distant NA2 isolates. Furthermore, two closely related Ontario isolates (W13-1 and W13-2) occupy intermediate positions between NA1 and the other North American samples, consistent with recent migration and admixture. Overall, these results indicate that while the NA1 lineage is strongly differentiated, the relationship between Ontario and the remaining North American isolates is less discrete and could potentially be explained by extensive sub-structure and isolation by distance within the NA2 lineage (see below).

**Figure 1:**
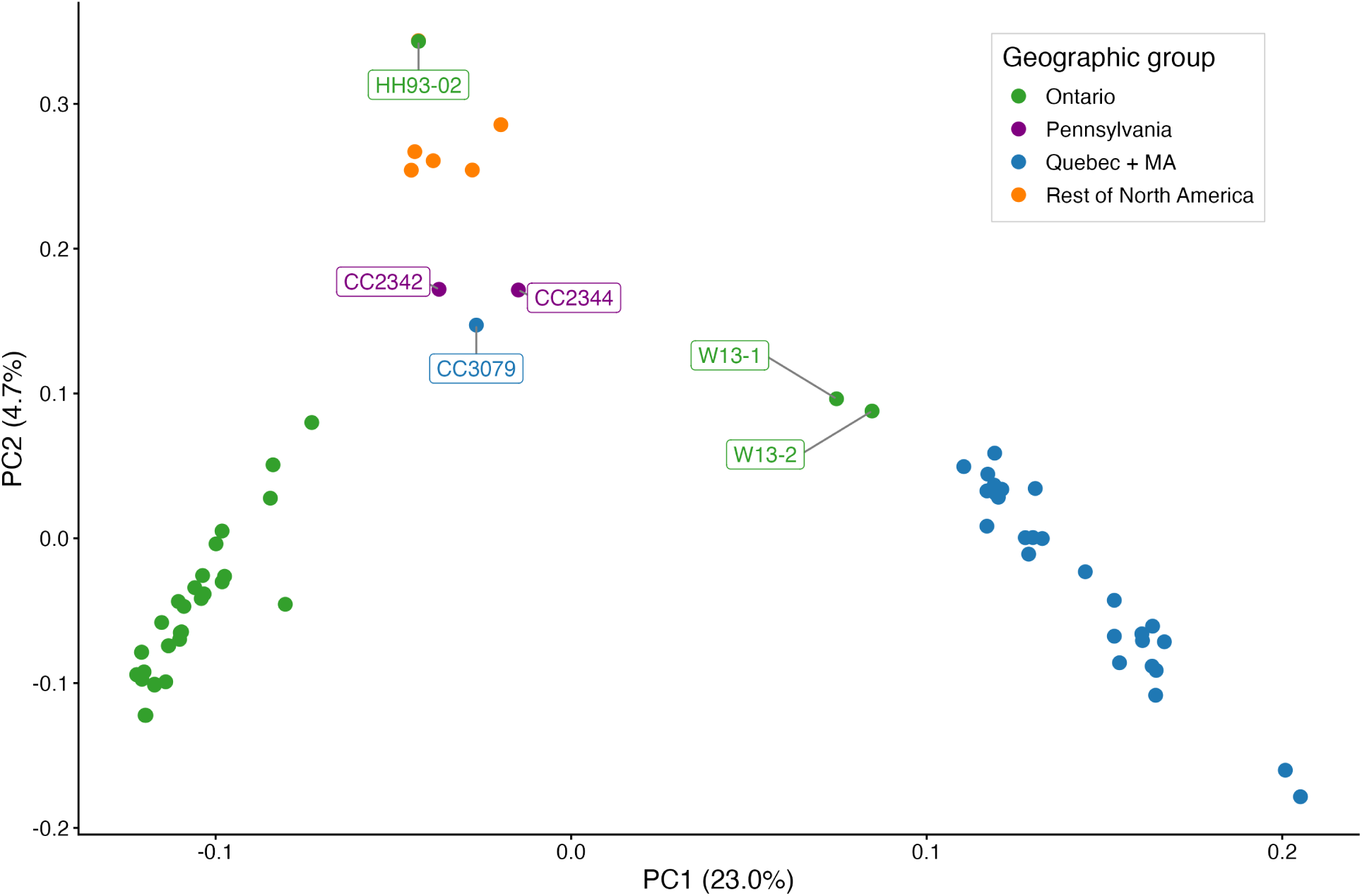
Population structure of North American *C. reinhardtii* isolates inferred by principal component analysis (PCA) based on genome-wide SNP variation. Points are coloured by geographic origin: Que-bec+Massachusetts (MA), Ontario, and the remaining North American locations. Four isolates (W13-1, W13-2, HH93-02, and CC-3079) are positioned in the PCA closer to samples from a different geographic group than their sampling origin and are labeled, consistent with migration or admixture.

Next, we evaluated population structure using STRUCTURE (testing K=1-10, 15 replicates each). Evanno’s ΔK statistic, obtained from STRUCTURE HARVESTER (Earl & vonHoldt, 2012), maximized at *K* = 2 (ΔK=2247.79), supporting a split where NA1 and Ontario are distinct, while NA2 is represented as an admixture between them (Figure S3). However, this method is known to be biased towards *K* = 2 even if further structure exists (Janes et al., 2017). Given the three distinct groups identified in our PCA and a secondary peak in Δ*K* at *K* = 3 (ΔK=13.45), we examined the *K* = 3 assignment plots to better resolve the relationships between known lineages and isolates sampled from Ontario (Figure 2a). This analysis was largely consistent with the three clusters defined in the PCA analysis, although all NA2 strains were recovered as highly admixed with the Ontario cluster, supporting a closer genetic relationship between these strains to the exclusion of NA1. Finally, fineSTRUCTURE analysis also revealed the same three clusters as PCA (Figure 2b), recovering both NA1, NA2, and an Ontario cluster. Consistent with the K=3 STRUCTURE analysis, NA2 and Ontario showed greater haplotype sharing with each other than with NA1, suggesting a more recent shared history or ongoing gene flow. As suggested by the PCA, the two Ontario isolates W13-1 and W13-2 were exceptions; STRUCTURE assigned ∼67-69% of their ancestry to NA1, with the remainder primarily to NA2, consistent with recent migration. All other Ontario isolates carried <10% NA1 ancestry, indicating limited gene flow from NA1 into the region. Conversely, the putative migrant Ontario strain HH93-02 exhibited strong haplotype sharing with the geographically diverse NA2 cluster. Overall, our results indicate three genetic clusters, with NA1 (Quebec+MA) strongly differentiated from the more closely related NA2 and Ontario clusters. However, this should be interpreted with caution and does not require that the Ontario-associated cluster represents a discrete, long-standing population. Both PCA- and fineSTRUCTURE-based methods are sensitive to uneven sampling (McVean, 2009; Puechmaille, 2016), and outside of Quebec and Ontario, most regions are represented by only one or two isolates. Under such conditions, densely sampled locations are more likely to appear as coherent clusters, whereas sparsely sampled regions may be artificially collapsed. Consistent with this, we previously showed that pairwise genetic distances among NA2 isolates follow a geographic pattern of isolation by distance (Craig et al., 2019). If similarly dense sampling were available across the geographic range of *C. reinhardtii*, what is currently labeled as NA2 may plausibly resolve into multiple localized populations or a genetic gradient across space that is only collectively distinguished relative to the divergent NA1 lineage. Accordingly, we treat the Ontario cluster as a well-sampled regional grouping whose relationship to NA2 remains unresolved, since it is plausible that it simply represents the only well-sampled population of NA2 rather than a fully independent population. We therefore use the labels “Quebec” and “Ontario” in their place to indicate that we have two densely sampled geographic regions that form independent genetic clusters. For all species-wide analyses, we consider all North American isolates.

**Figure 2:**
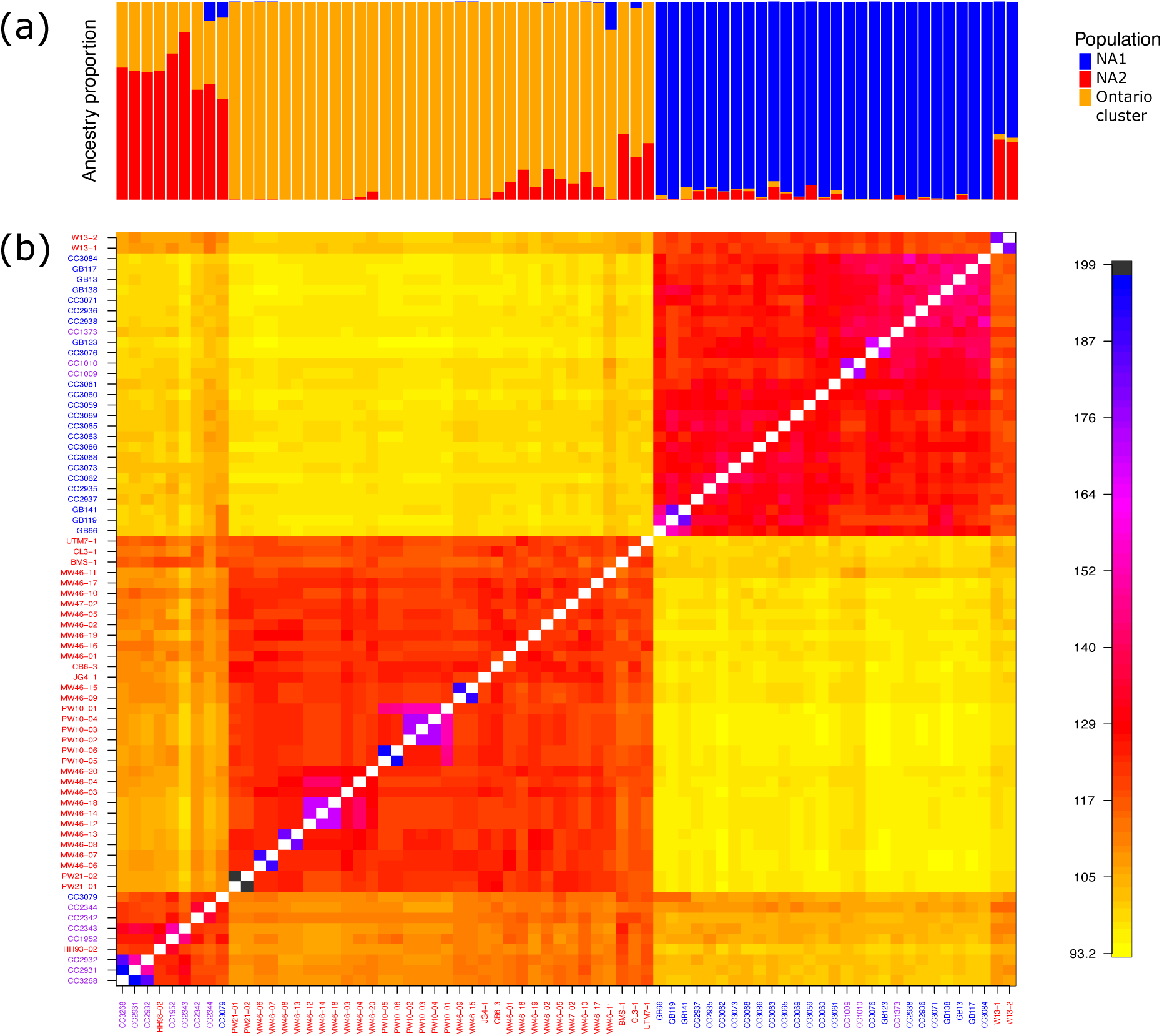
(a) STRUCTURE plot of isolates for K=3. Colour indicates ancestry proportion to a specific cluster. (b) Coancestry matrix of 73 isolates of *C. reinhardtii*. The colour of any cell at position (i, j) in the coancestry matrix M indicates the number of haplotypes shared between individuals i (as recipient) and j (as donor), where M[i, j] denotes the number of shared haplotypes.

### 3.2 Clonality and sampling diversity

We assessed whether our new isolates included genetically distinct isolates or clones of the same genotype. We identified eight clonal groups comprising 18 isolates (Table S5). Most groups consisted of pairs, although two groups contained three isolates (PW10-2/3/4 and MW46-12/14/18). After selecting one representative of each clonal group, 24 unique genotypes remained in the new samples we report here, or 31 including the seven Ontario genotypes reported recently (Ford et al., 2022). No clonal group spanned multiple sampling locations, indicating no widespread or highly abundant genotypes across the landscape. However, of these clonal groups, seven included isolates recovered from different soil samples at the same location (PW10-2/3/4, PW10-5/6, PW21-1/2, MW46-8/13, MW46-9/15, MW46-12/14/18, and HH93-1/2), demonstrating that these are free-living clonal individuals (biological clones) as opposed to clones that arose during lab cultivation (technical clones). Two out of the five *C. reinhardtii*-positive sites yielded more than one unique genotype, including one instance where genetically distinct strains were isolated from a single soil sample (MW46-12/14 and MW46-13). Together, these results indicate that multiple genotypes may often coexist at fine spatial scales, while landscape-scale clonality is absent from our current sampling. In addition to clear clonal sets, our analysis revealed several pairs of isolates with exceptionally high haplotype sharing that likely represent closely related individuals, rather than identical clones, as is the case for the “laboratory strains” of *C. reinhardtii*, represented herein by CC-1009 and CC-1010 (Gallaher et al., 2015). These relationships are consistent with local sexual reproduction but were not treated as clonality in our analyses. The putative admixed strains W13-1 and W13-2 (Figure 1) fall within this category, suggesting that they may be descendants of a single migration event.

### 3.3 Whole-genome and regional genomic diversity, divergence, and differentiation

We quantified within-population diversity (*π*), nucleotide divergence (d*_xy_*), and differentiation (F*_st_*) using pixy (Korunes & Samuk, 2021) (Figure 3). Considering all strains from each geographic location, genome-wide *π* was slightly higher in Quebec (0.02487) than in Ontario (0.02266), and the two locations were highly differentiated (d*xy* = 0.03384, F*st* = 0.2764). When considering broader population structure (i.e. accounting for clustering of putative migrant or admixed strains), diversity varied among the three defined clusters. Consistent with its broad geographic range, NA2 exhibited the highest diversity (*π* = 0.0317), followed by NA1 (*π* = 0.0246), while the Ontario-associated cluster had somewhat lower diversity (*π* = 0.0199) (Figure 3a). Patterns of divergence and differentiation were consistent with results from the population structure analyses. Strains associated with the Ontario cluster exhibited less differentiation from NA2 (d*xy* = 0.0296, F*st* = 0.0800) but were highly differentiated relative to NA1 (d*xy* = 0.0343, F*st* = 0.3372), while NA1 and NA2 were also divergent (d*xy* = 0.0355, F*st* = 0.1755) (Figure 3b, c).

**Figure 3:**
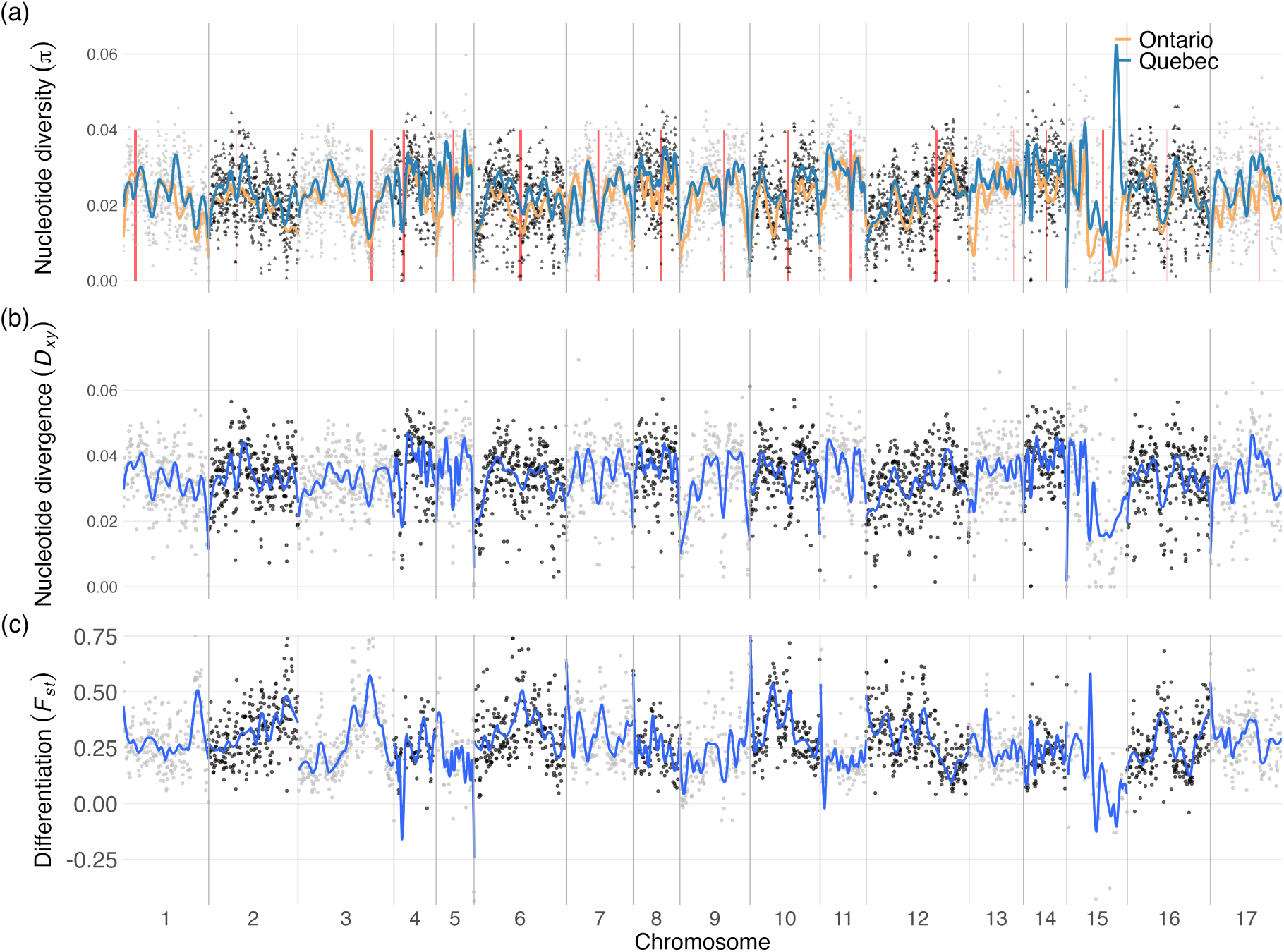
From top to bottom, panels show genomic landscape of (a) nucleotide diversity, *π* (b) nucleotide divergence, d*_xy_* and (c) differentiation, F*_st_* across the *C. reinhardtii* genome in non-overlapping 30 kb windows. d*_xy_* and F*_st_* were both calculated between Ontario and Quebec. Red vertical bars indicate centromere positions.

Genome-wide diversity was correlated between Ontario and Quebec across matched genomic windows (Pearson’s *r* = 0.589). Previous analyses of genome-wide variation in diversity were performed on earlier versions of the reference genome, which featured a number of inter-chromosomal misassemblies and lacked mapped centromere locations (Craig et al., 2022). We found that nucleotide diversity was significantly reduced in centromere-proximal regions relative to chromosome arms. In Quebec, diversity near centromere was reduced by 8.74% (Δ*π* = 0.00222, p = 0.0013). In Ontario, the effect was in the same direction but weaker at 5.26% (Δ*π* = 0.00119, p = 0.044). These regions were defined as 1 Mb windows centered on the midpoint of centromeres (which have a mean length <200 kb (Craig et al., 2022)), and thus the reduction in diversity is not expected to be an artefact of low mapping quality in these repetitive regions. Diversity was even more strongly suppressed in subtelomeric regions: median *π* in the terminal 50 kb with a 56.61% proportional reduction compared to chromosome arms in Ontario (0.0098 vs. 0.0225; Wilcoxon rank-sum test, *p* = 5.7 *×* 10*^−^*^21^) and 54.88% in Quebec (0.0113 vs. 0.0250; *p* = 2.7 *×* 10*^−^*^25^). Thus, many broad scale factors shaping genomic variation in genetic diversity may be shared across the species, with the most pronounced diversity reductions occurring in centromere-proximal regions and at chromosome ends. Reduced diversity at chromosome ends is consistent with previous work showing decreased nucleotide diversity and elevated linkage disequilibrium near chromosome termini in *C. reinhardtii* (Flowers et al., 2015), supporting the conclusions that selection at linked sites has a widespread influence on genomic diversity (Hasan & Ness, 2020). However, the apparent reductions near centromere-proximal regions have not yet been explicitly examined in relation to the recombination landscape and may represent an additional genomic feature shaping diversity in this species.

### 3.4 Species-wide and population-specific selective sweeps

To detect species-wide patterns of selection, we performed selective sweep analyses on all samples together. We first used background selection simulations to generate null distributions of the composite (*µ*) statistic under the inferred demographic model. Model fitting indicated that a constant population size best matched the observed site frequency spectrum, and additional historical events (such as bottlenecks or admixture) did not improve fit. We therefore simulated *µ* values under this minimal model to determine a threshold value to identify outliers. Given that the maximum simulated *µ* value should represent a conservative cutoff, we expected this threshold to identify the strongest empirical outliers. Instead, it classified 4,020 genes (∼24% of all genes) as candidates, suggesting that the background-selection simulations produced a null distribution whose upper tail was too low relative to the empirical data. We therefore treated this simulation-derived cutoff as overly permissive in practice and instead used the top 0.1% of genome-wide *µ* values as a more stringent empirical threshold (Figure 4a). With this approach we found 252 distinct regions including 893 genes that intersected putative sweep windows.

**Figure 4:**
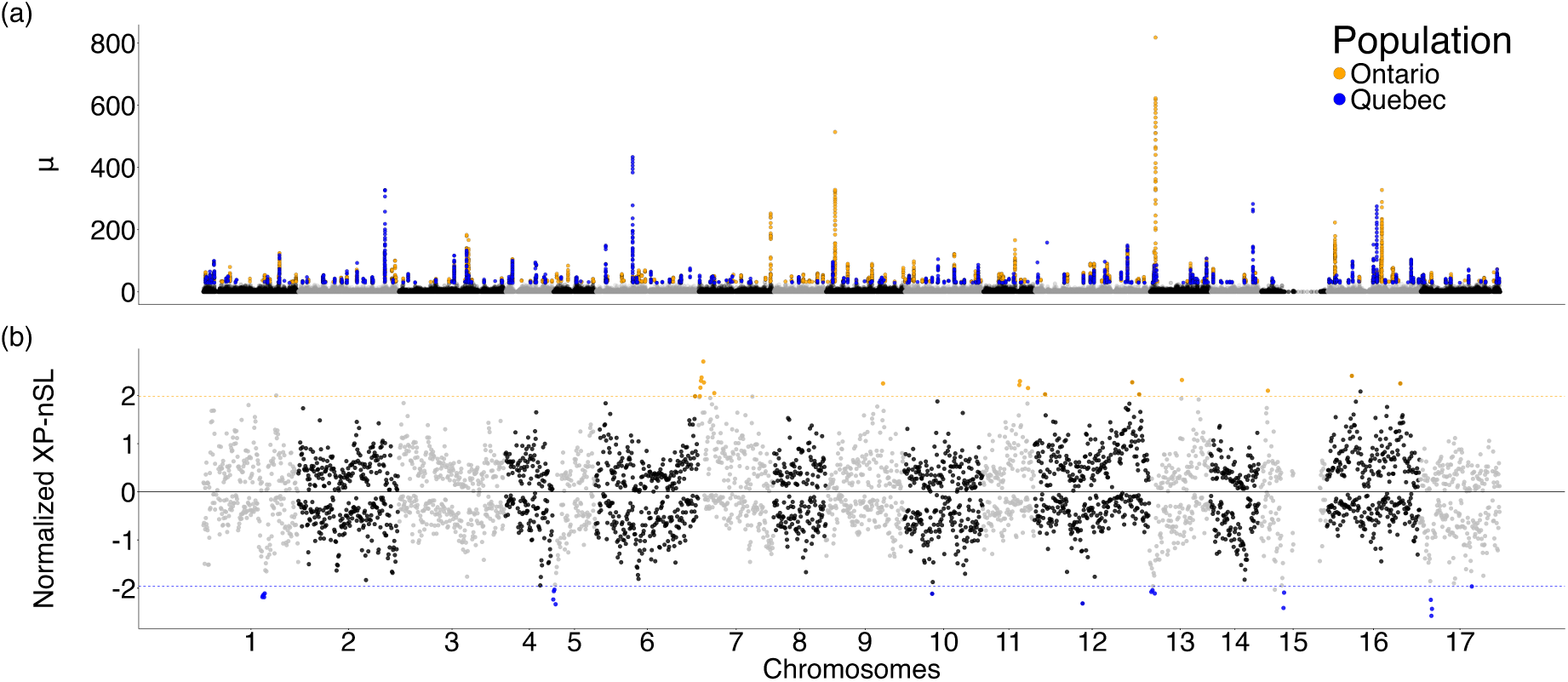
Manhattan plot of the chromosome-by-chromosome landscape of selective sweep scores in the *C. reinhardtii* genome from *µ* and normalized XP-nSL values. (a) *µ* values calculated across the genome for Ontario (orange) and Quebec (blue) clusters, with top 0.1% outlier windows highlighted in their respective population colours. (b) Normalized XP-nSL values calculated in 50 kb windows contrasting Ontario and Quebec. Higher positive values indicate stronger sweep signals in Quebec, whereas more negative values indicate stronger signals in Ontario. Outlier windows are highlighted in orange (Ontario) and blue (Quebec).

Approximately 5.09% of the genome (5.72 Mb) falls within these windows species-wide. After we merge adjacent sweeps, we find that the mean length of a sweep is 22.7 kb wide and harbours a median of three genes, although the modal number of genes in a window is one. To browse each of the genes, see Table S7. *µ* values show an exponential decline (Figure S5), indicating that a small subset of genomic regions experience strong selection. After identifying the top 0.1% outlier windows, we tested whether sweeps clustered on specific chromosomes. A length-corrected *χ*^2^ test found no evidence of chromosomal enrichment, indicating that sweeps are broadly distributed.

We next investigated population-specific patterns of selection using Quebec and Ontario isolates separately, while excluding recently admixed individuals (see Methods). Using the same approach as above we revealed that *µ* shows a similar exponential decline in both population samples (Figure S5). Ontario displays a markedly longer right tail, with maximum values 15.5% higher than Quebec. Although we found 223 and 189 sweeps in Ontario and Quebec, these sweep candidates also included shared sweeps that are species-wide. When the species-wide regions are excluded, we are left with 168 and 133 population-specific sweep signatures in Quebec and Ontario. These regions cover 2.92% and 1.61% of the genome in Quebec and Ontario respectively.

To better examine population-restricted sweeps we employed the XP-nSL statistic which identifies genome regions where one population has relatively longer haplotype blocks that are indicative of recent positive selection (Szpiech et al., 2021). With this approach we found 39 candidate sweeps, with 19 and 20 in Ontario and Quebec, respectively (Figure 4b). In line with expectations, outlier Quebec *µ* regions had a significant overlap with outlier Quebec XP-nSL regions (3 of the 20 XP-nSL windows, p=0.005), and outlier Ontario *µ* regions had a significant overlap with the outlier Ontario XP-nSL regions (9 of the 19 XP-nSL windows, p=0.001).

We also found that six of the 39 population restricted XP-nSL windows overlapped with species-wide sweeps identified from *µ* values. This indicates that the sweep signal in one population may be strong enough to create a species-wide signal despite selection in only one population. Supporting this view, the six loci had no evidence of a sweep in the population where no selection was expected when considering population-specific *µ* values.

In sum, we found that there was evidence of species-wide selective sweeps at 151 loci, overlapping 519 genes. We also find that there are a total of 325 regions that appear to have exclusively swept in one population, with 179 and 146 in Ontario and Quebec respectively. It is worth noting that using statistical thresholds to identify swept regions means that the exact numbers here will vary depending on the stringency of the threshold. Considering figure Figure 4 it is evident that there are a small number of very pronounced peaks where evidence of a sweep is much higher than the threshold. These observations demonstrate that adaptive evolution occurs both across the species range and along population-specific trajectories.

### 3.5 Species-wide targets of selection

Examination of broad-scale patterns of the functions of genes in selected windows can provide insights into adaptation of microbes, where ecological and phenotypic observations are challenging. GO enrichment analysis revealed that there were species-wide signatures of selection in two broad areas, detoxification of methylglyoxal and viral defense pathways (Table S8). Active selection on immunity is consistent with long-standing patterns that immune genes are frequent targets of positive selection (Bonhomme et al., 2015; Hu et al., 2022). The enrichment for immunity related functions is partly driven by a compelling species-wide candidate sweep that was a ∼50 kb region on chromosome 13 (position ∼4.9 Mb) that contained eight genes. There is a cluster of five homologous genes in this region with GO terms are associated with immunity functions, specifically double-stranded RNA binding (for full information, see Table S9). Notably, these genes contain a predicted oligoadenylate synthase (OAS) domain, which is present in a key antiviral effector in animals that detects dsRNA to initiate viral RNA degradation (Schwartz & Conn, 2019). A recent survey of eukaryotic innate immunity identified OAS-like CD-NTase homologs within the Archaeplastida and suggested they represent an ancient inheritance from the earliest eukaryotes (Culbertson & Levin, 2023). However, the evolutionary history and functional role of the OAS-like genes in this *C. reinhardtii* region remain unresolved and will require targeted phylogenetic and experimental analyses. We therefore interpret this locus as a strong candidate immune-related sweep region, illustrating how population genomic scans can nominate previously uncharacterized gene clusters for future functional study in green algae.

### 3.6 Population-specific targets of selection

To identify individual genes with strong signals of having undergone selection in one population to the exclusion of the other, we identified areas in the genome with clusters of significant XP-nSL values (measured in 5 kb windows) coincided with regions where differentiation (F*_st_*) values approached 1 (>= 0.95). We additionally performed manual inspection of nucleotide diversity (*π*) across these regions, flagging loci where the two populations showed marked divergence in *π* relative to the surrounding genomic background. For all predicted functional annotations identified using this method, see Table S10. We identified 33 genes across eight genomic regions that represent strong candidates for local adaptation, resolved into two distinct sets corresponding to the Quebec and Ontario populations. A sweep region on chromosome 1 (Ontario) remains one of the most pronounced signals genome-wide. This interval contains two functionally uncharacterized genes, a voltage-gated chloride channel adjacent to a predicted calcium/calmodulin-regulated kinase, and shows coordinated evidence of recent selection: *π* indicates a localized collapse of diversity in only the focal population, XP-nSL values exceed significance thresholds, and sweep windows exhibit *F_st_* values approaching 1.0 (Figure 4; Figure 5). Beyond this focal case, similar functions appear among Ontario candidates on chromosome 17, which include a voltage-gated Ca^2+^ channel (*CAV5*) and two GPR1/FUN34/YaaH family transporters (*GFY4* and *GFY5*), putative membrane proteins implicated in small-molecule transport, including acetate (Durante et al., 2019), suggesting that ion handling and small-molecule transport may be recurring targets of population-specific adaptation in Ontario. By comparison, Quebec sweeps predominantly contain genes associated with transcriptional regulation and RNA metabolism. On chromosome 7, Quebec candidates include DEAD-box RNA helicase, RNA polymerase Rpc34 subunit family protein, and a nearby Chlamydomonas-specific G-protein. Across both Quebec and Ontario, a smaller number of genes fall into general metabolic or proteostasis-related functions (e.g., aspartyl aminopeptidase and aspartyl protease). Together, these patterns indicate that both populations experienced distinct adaptive trajectories.

**Figure 5:**
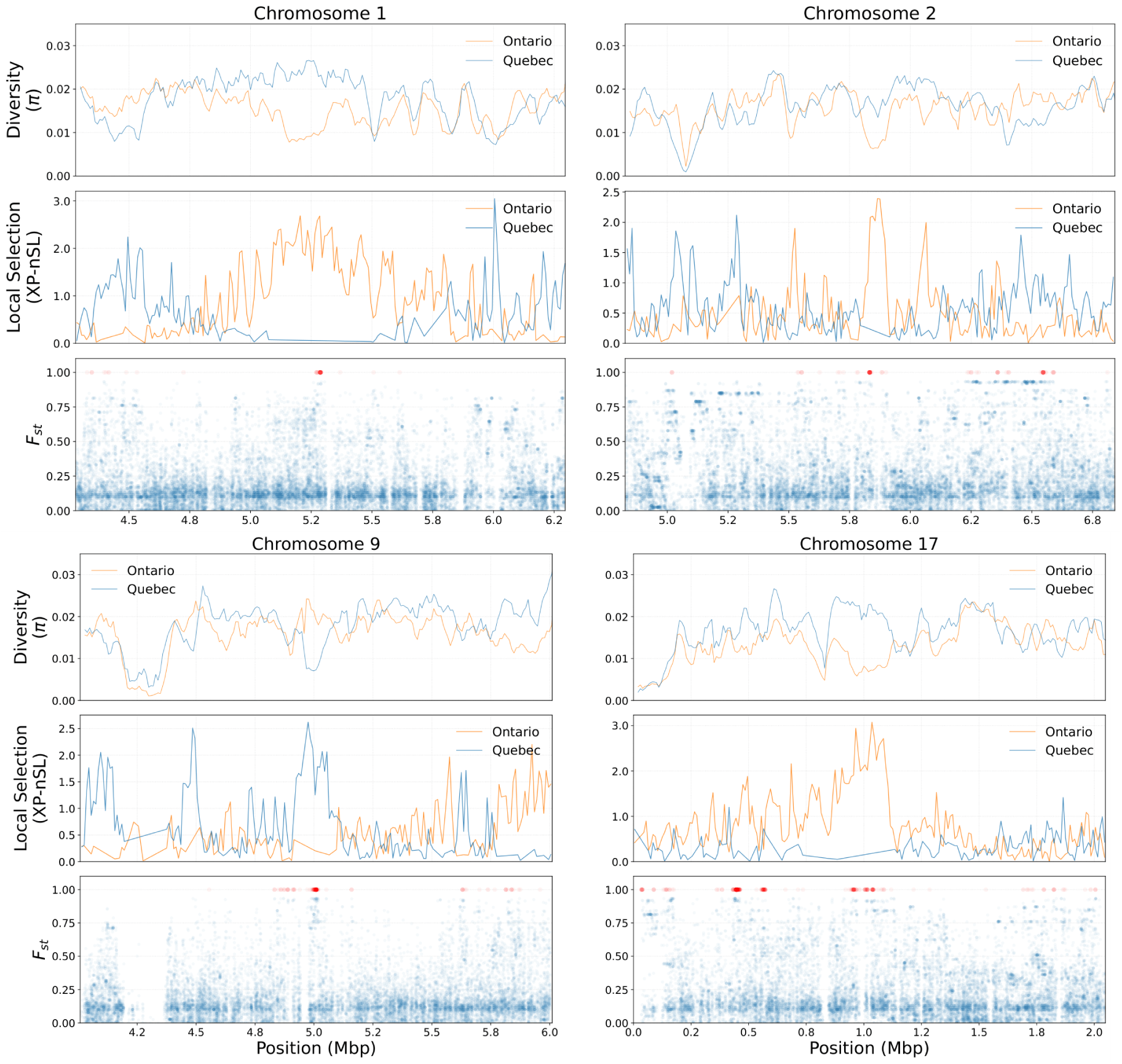
Array of genomic signals supporting examples of selective sweep on chromosomes 1, 2, 9, and 17 in *C. reinhardtii*. Each chromosome-specific panel uses a shared genomic coordinate axis and contains three tracks. From top to bottom: (i) windowed estimates of nucleotide diversity (*π*) across the interval for Ontario isolates (blue) and Quebec isolates (orange); (ii) local selection (XP-nSL) values for Ontario (blue) and Quebec (orange), absolute values taken of Ontario XP-nSL values; (iii) per-window *F_st_* between Ontario and Quebec isolates, with all windows shown in blue and those with extreme differentiation (*F_st_≥* 0.95) highlighted in red.

## 4 Discussion

In this study we have characterized the genomic diversity of 38 new isolates of *C. reinhardtii* from Ontario including seven our group recently reported from the same region (Ford et al., 2022). These new isolates represent both a novel geographic population and a doubling of the known sampling of *C. reinhardtii*, facilitating intra- and inter-population microbial population genetic exploration. These samples also provided us insight into the clonal structure of *C. reinhardtii* in nature. Interestingly, identified clones were never from different sampling locations, meaning that we found no evidence of widespread clones. In fact, in two out of five newly reported sampling sites, we found distinct genotypes from the same site. In one case, a single sampling tube had contained genetically distinct strains, providing evidence for fine-scale local diversity. Such fine-scale co-occurrence of diverse strains is consistent with ecological conditions that facilitate sexual reproduction, where successful mating requires close spatial proximity of compatible genotypes. Our new sampling of *C. reinhardtii* enables novel insights into microbial diversity, migration patterns, and history of selection.

The genetic diversity of *C. reinhardtii*, has previously been shown to be relatively high (Craig et al., 2019; Flowers et al., 2015). Because of our expanded sampling of the species and a new version of the reference genome, we reestimated diversity values for the previously defined and new clusters. We found that diversity among Ontario and Quebec strains was high relative to many other species (Leffler et al., 2012), likely reflecting a large effective population size of *C. reinhardtii* in these regions. Interestingly, when investigating population genetic structure, Ontario strains are primarily grouped into a new cluster with lower diversity (*π* = 0.0199) than NA1 and NA2 respectively (0.0246, 0.0317). The high diversity of NA2 likely reflects that its sampling is geographically dispersed and may indicate that more dense sampling in the USA would uncover several distinct population clusters that fall within the NA2 lineage. Lower divergence and differentiation values were observed between NA2 and Ontario (d*_xy_* = 0.0296, F*_st_* = 0.0800) compared to either NA1 vs Ontario (d*_xy_* = 0.0343, F*_st_* = 0.3372) or NA1 and NA2 (d*_xy_* = 0.0355, F*_st_* = 0.1755). These relationships reflect either shared ancestry or ongoing gene flow between NA2 and Ontario, and as discussed previously it may be the case that Ontario simply represents a densely sampled population from within a broad NA2 assemblage. Overall, the patterns of structure we see are consistent with a model of allopatric divergence in glacial refugia, as proposed for *C. reinhardtii* and other microbial eukaryotes. In particular, the observed pattern mirrors that described in wild yeast (*Saccharomyces paradoxus*), where divergence between northeastern and more broadly distributed North American lineages has been attributed to isolation in the Atlantic and Mississippian refugia during the last glacial period (Charron et al., 2014). Under this framework, NA1 likely derives from an Atlantic refugium east of the Appalachians, whereas all other samples reflect expansion from more southern and western refugia (Craig et al., 2019). In line with this, both the Ontario and originally defined NA2 strains are clearly highly differentiated from the Quebec and Massachusetts NA1 strains. Attempting to sample new isolates from potential contact zones towards the St. Lawrence River would be an interesting future direction.

Levels of diversity along the genome were correlated between Ontario and Quebec (Pearson *r* = 0.59), consistent with shared ancestry and landscape of background selection. However, substantial population-specific variation remains (approximately 65% of variance in *π*), indicating that demographic differences, local selection, mutation, or drift have also shaped diversity independently in each group. Under linked selection, nucleotide diversity is expected to be reduced in genomic regions where recombination is suppressed, because selection affects linked neutral sites over broader physical distances (Begun & Aquadro, 1992). Consistent with this expectation, we observed significantly reduced nucleotide diversity near centromeres in both Ontario and Quebec, with a stronger reduction in Quebec. While a positive relationship between recombination rate and nucleotide diversity has previously been reported in *C. reinhardtii* (Hasan & Ness, 2020), earlier analyses relied on fragmented genome assemblies that did not allow precise localization of centromeress. Notably, diversity reduction was even stronger in subtelomeric regions, where median *π* is reduced by over 50% within 50 kb of chromosome ends. This pattern is consistent with reduced recombination near telomeres (Flowers et al., 2015). Together, these patterns of diversity highlight that variation in recombination is an important determinant of diversity in the *C. reinhardtii* genome.

A core prediction of the ubiquity model is that migration among populations should be widespread. Previous work had already found that NA1 strains showed signatures of admixture with NA2 (Craig et al., 2019). Here, our new sampling provides clear evidence that *C. reinhardtii* exhibits substantial levels of migration. Three of the Ontario isolates clustered more closely with those from other geographic regions, including HH93-02, which clustered with NA2, and W13-1 and W13-2, which appear to be NA1-NA2 hybrids. These add to a previously identified migrant, CC-3079, which appears to have migrated from NA2 to Quebec (Craig et al., 2019). Together this means 6.3% of 63 unique North American strains are migrants, and such a high level of migration should lead to homogenization of genetic structure, if there is no impediment to introgression (Sundqvist et al., 2016). The observation of regular migration falls in line with the prediction of the ubiquity model, that microbes frequently disperse over large geographic distances. This raises the question as to why the present genetic differentiation between lineages has been maintained since the glacial retreat. While it is possible that high migration is a recent anthropogenic phenomenon, or that reproductive isolation is maintaining population structure, local adaptation may impede the introgression of migrant DNA. To examine the role played by selection in explaining our data, we examined signatures of genome-wide selective sweeps across the range.

If population structure was primarily maintained through geographic isolation, as predicted by the endemicity model, then species-wide selective sweeps should be uncommon, as the lack of gene flow between populations would impede the spread of new beneficial alleles. To the contrary, we find that species-wide sweeps are quite common, identifying 151 non-overlapping sweep regions. The presence of so many sweeps that affect all samples provides evidence for selection across *C. reinhardtii*. This result is only consistent with the endemicity model if these sweeps either predate the split into current subpopulations or if, selection acted repeatedly on ancestral variation in each subpopulation. To examine if the species-wide sweeps predate the split in the common ancestor, we compared reductions in diversity in species-wide sweeps (81.9%) to reductions in each of the population-exclusive sweeps (between 77.4% and 90.2%). Seeing that these are comparable, we find that it is unlikely that most species-wide sweeps are substantially older, although we can not rule out the possibility that beneficial alleles were inherited from the ancestor but underwent sweeps in both subpopulations at a later period.

We hypothesize that selection contributes to the maintenance of population subdivision in *C. reinhardtii*, and that this should be identifiable in differences between the genomic targets of sweeps across populations. Combining our population-specific sweep scans we found a total of 146 Ontario-specific sweeps and 179 Quebec-specific sweeps. Local selection in combination with the existence of early generation migrants and population structure is consistent with the ubiquity model’s predictions that these microbes are capable of movement but selection in the local environment poses some barriers to survival and therefore complete introgression. This has previously been seen in the cosmopolitan microbial eukaryote and marine diatom, *Thalassiosira rotula*, which exhibits near-panmictic gene flow across ocean basins and negligible isolation by distance, yet maintains distinct genetic clusters aligning with environmental and ecological gradients (Whittaker & Rynearson, 2017). However, it remains possible that other forms of cryptic reproductive isolation are operating to maintain population structure in *C. reinhardtii*, such as mating incompatibilities between subpopulations or temporal segregation of sexual reproduction driven by environmental or seasonal differences (R. Ji et al., 2010). The strongest test of local adaptation would require reciprocal transplants which are challenging in a soil dwelling microbe. Alternatively, candidate phenotypes that might drive local adaptation could be proposed from the genes in these sweeps, which we explore further below.

The expectation from a selective sweep is that distortions of genetic diversity and the allele frequency distribution as well as linkage should dissipate over the genomic distance from the selected locus. Therefore, the genes or loci subject to selection can be inferred to be those central to a given sweep signature. The sweep windows in *C. reinhardtii* are relatively narrow (mean 22.7 kb) and most often contain a single gene. These windows are relatively narrow compared to many other species where selective sweeps can span more than 10^5^–10^6^ bp (Clark et al., 2004; Voight et al., 2006), and this difference might reflect a high population recombination rate facilitated by the large *N_e_*of *Chlamydomonas* (Hasan et al., 2019). The functional composition of lineage-specific sweep-associated genes, based on the combined evidence of extreme XP-nSL and high differentiation, differed between Ontario and Quebec. Ontario candidate loci were enriched for genes involved in membrane transport, including multiple voltage-gated channels and GPR1/FUN34 transporters, as well as proteolysis, oxidative stress response, and carbohydrate metabolism. By contrast, Quebec candidate loci were dominated by genes associated with transcription and RNA processing. These contrasts should be interpreted cautiously because the exact causal loci within swept regions cannot be verified from sweep scans alone. Nevertheless, the non-overlapping functions associated with swept regions in each population suggest that Ontario and Quebec may be responding to different ecological pressures. Such population-specific selection could contribute to barriers to gene flow if migrants or hybrids carry alleles that are maladapted to local environmental conditions. Altogether, the functional targets of selection reveal both shared selective pressures across North America and population-specific ecological challenges, offering potential mechanisms by which selection can impede the homogenizing influence of ongoing migration.

## 5 Conclusion

Our study represents a substantial advance in sampling the model green alga *C. reinhardtii*, more than doubling the number of known strains. Enhanced geographic coverage enabled us to gain novel insights into fine-scale clonal diversity and the detection of three strains that represent migrants between geographically distant locations. The maintenance of distinct lineages appears paradoxical when considering that roughly 6% of characterized strains show evidence of recent dispersal. This phenomenon—genetic subdivision coexisting with active dispersal—has been documented in other free-living microbial eukaryotes (Whittaker & Rynearson, 2017). Our hypothesis is that endemicity can occur when there is true allopatry, but in the presence of microbial migration, environmentally-mediated selection contributes to the preservation of differentiation. The role of adaptive evolution in structuring microbial populations has some prior empirical support. Specifically, selective sweeps in *Neurospora crassa* revealed two genomic islands under divergent selection between populations that moderated the proliferation of this fungus in environments of differing temperatures. Our results show that *C. reinhardtii* clearly possesses the dispersal capacity predicted by ubiquity theory, as multiple migration events and admixed genotypes demonstrate. However, proving equally important, divergent selective pressures on the environmental population of endemic populations may counteract the homogenizing effects of gene flow. Rather than supporting one model over the other, these findings suggest that while allopatry in the past was a dominant force shaping microbial biogeography there is clear evidence of population-specific adaptation that may maintain structure despite ongoing dispersal.

## 6 Supplementary

**Figure S1:**
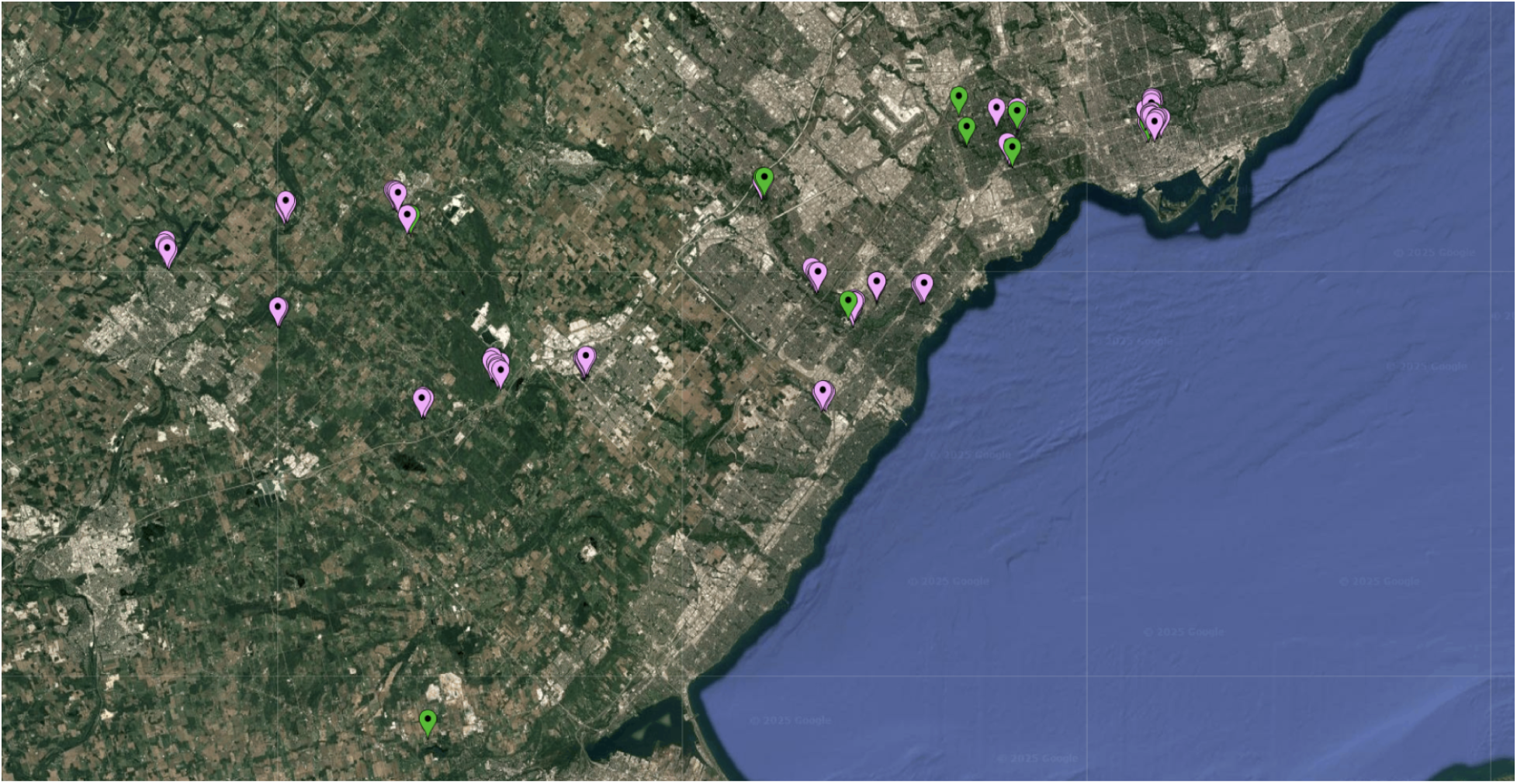
Map of sampling locations surveyed for *Chlamydomonas reinhardtii* in southern Ontario. Green markers indicate sites yielding one or more *C. reinhardtii* isolates, whereas purple markers indicate surveyed sites where *C. reinhardtii* was not detected. The spatial distribution highlights both the patchy occurrence of *C. reinhardtii* and the extensive geographic coverage of the sampling effort.

**Figure S2:**
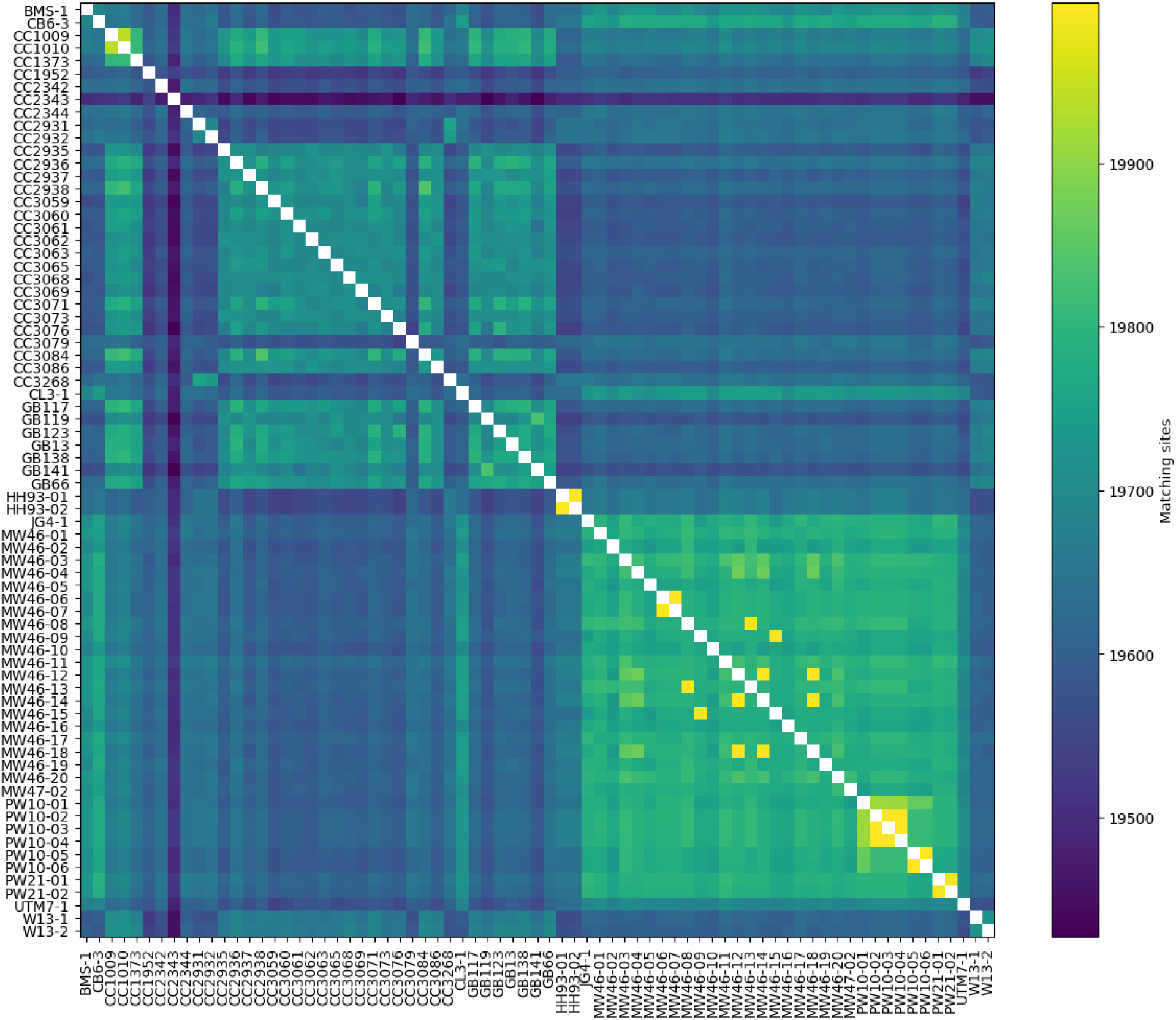
Pairwise genomic similarity among *Chlamydomonas reinhardtii* samples. Each cell represents the number of matching sites observed between a pair of strains, estimated from 20,000 randomly sampled genomic positions. Colour intensity indicates the strength of similarity, with lighter colors corresponding to a greater number of matching sites. A small number of strain pairs exhibit near-complete identity, forming distinct high-similarity blocks consistent with clonal relationships, while the majority of comparisons show substantially lower similarity, reflecting non-clonal divergence.

**Figure S3:**
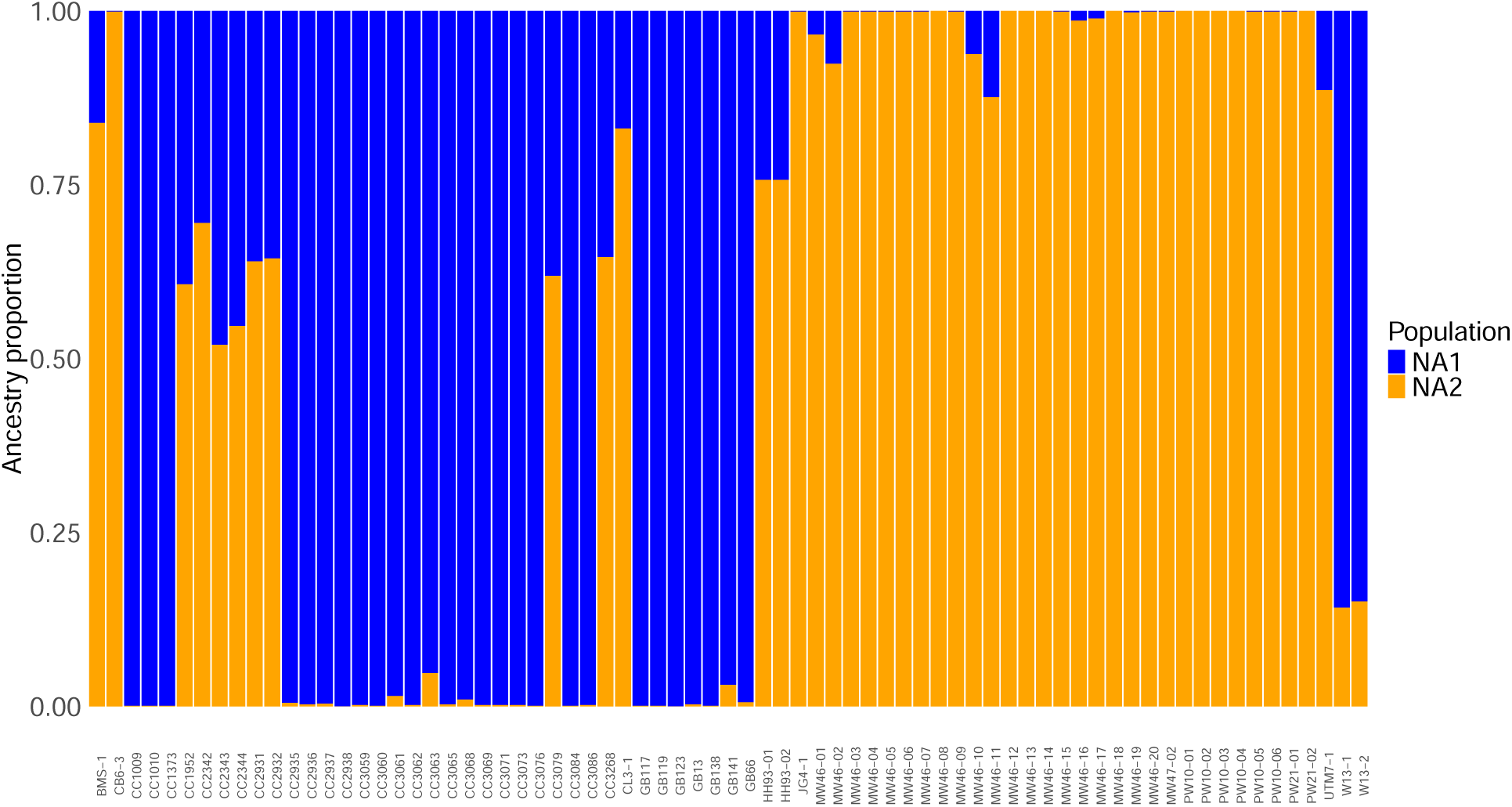
STRUCTURE PLOT of isolates when K=3. Colour indicates ancestry proportion to a specific cluster.

**Figure S4:**
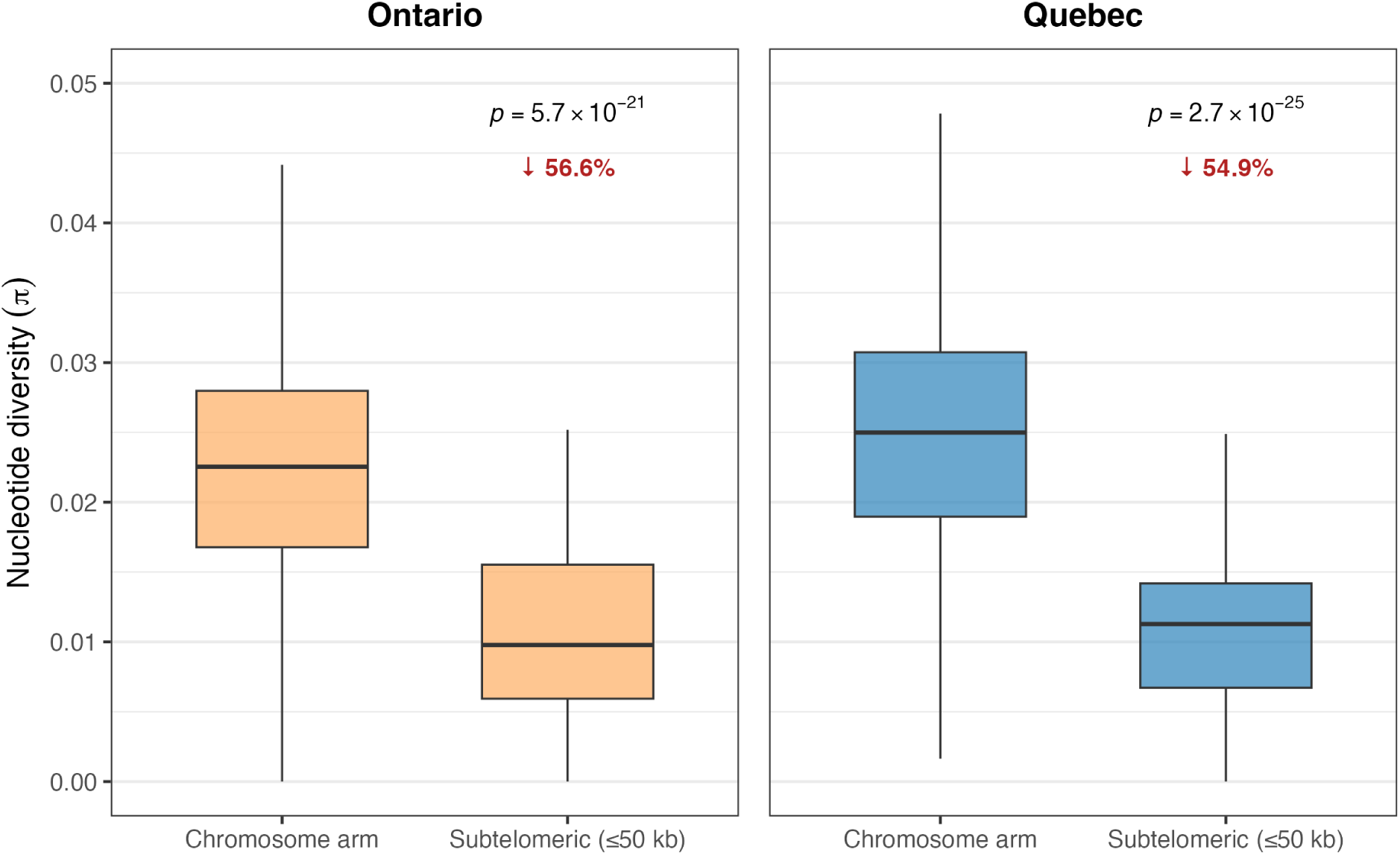
Subtelomeric suppression of nucleotide diversity in *Chlamydomonas reinhardtii*. Boxplots compare the distribution of *π* in chromosome arms versus subtelomeric regions (*≤*50 kb from chromosome ends) for Ontario (left) and Quebec (right) populations. Individual genomic windows are shown as semi-transparent points (subsampled for visibility). Median *π* was reduced by 56.6% in Ontario (Wilcoxon rank-sum test, *p* = 5.7 *×* 10*^−^*^21^) and 54.9% in Quebec (*p* = 2.7 *×* 10*^−^*^25^), consistent with strong linked selection in regions of suppressed recombination near telomeres.

**Figure S5:**
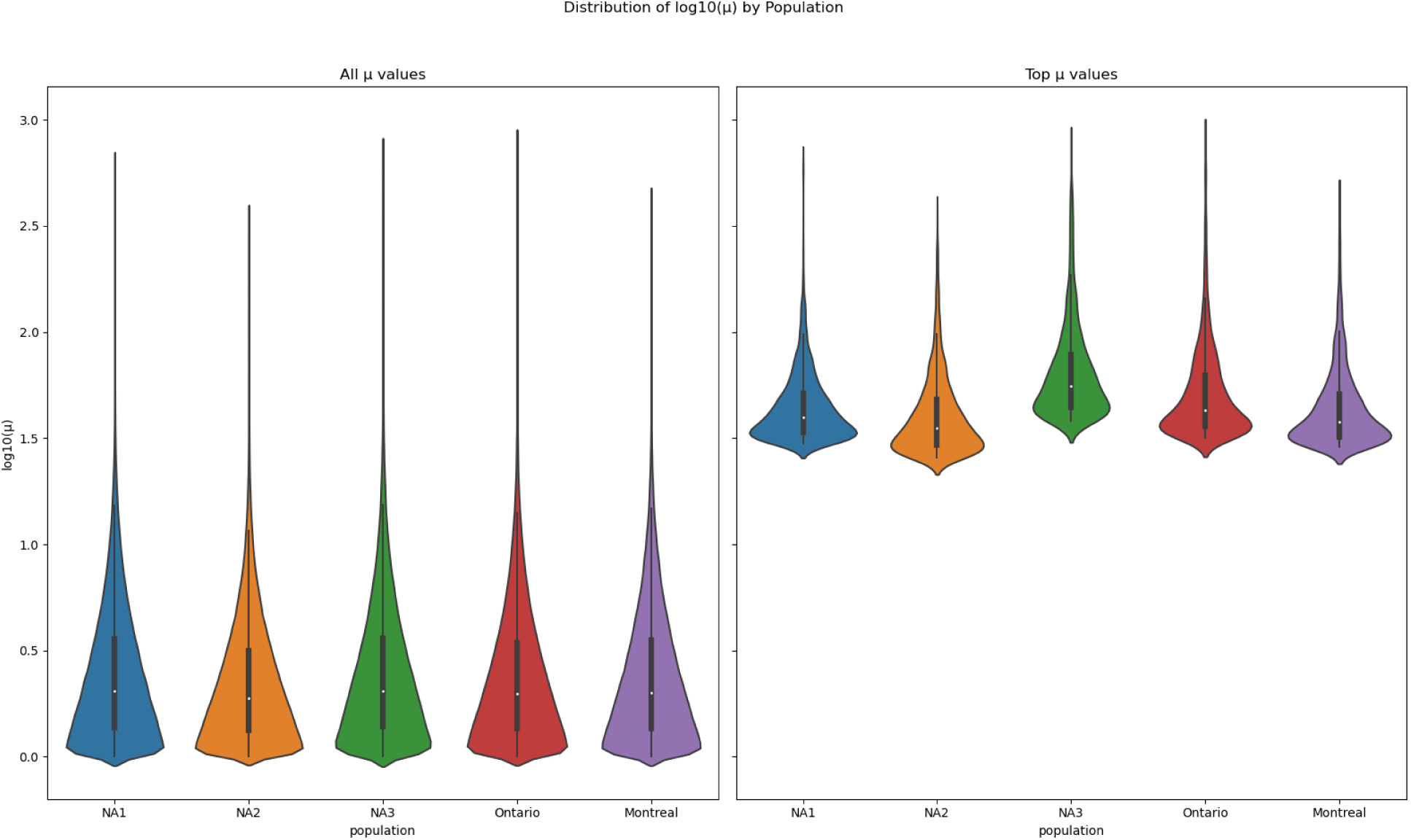
Distribution of *µ* values per population. Top *µ* values are defined as those exceeding the cutoff for being among the most 0.1% extreme values.

## References

Abondio, P., Cilli, E., & Luiselli, D. (2022). Inferring Signatures of Positive Selection in Whole-Genome Sequencing Data: An Overview of Haplotype-Based Methods. Genes, 13 (5), 926. 10.3390/genes13050926

Ailloud, F., Didelot, X., Woltemate, S., Pfaffinger, G., Overmann, J., Bader, R. C., Schulz, C., Malfertheiner, P., & Suerbaum, S. (2019). Within-host evolution of Helicobacter pylori shaped by niche-specific adaptation, intragastric migrations and selective sweeps. Nature Communications, 10 (1), 2273. 10.1038/s41467-019-10050-1

Alachiotis, N., & Pavlidis, P. (2018). RAiSD detects positive selection based on multiple signatures of a selective sweep and SNP vectors. Communications Biology, 1 (1), 1–11. 10.1038/s42003-018-0085-8

Badouin, H., Gladieux, P., Gouzy, J., Siguenza, S., Aguileta, G., Snirc, A., Le Prieur, S., Jeziorski, C., Branca, A., & Giraud, T. (2017). Widespread selective sweeps throughout the genome of model plant pathogenic fungi and identification of effector candidates. Molecular Ecology, 26 (7), 2041–2062. 10.1111/mec.13976

Begun, D. J., & Aquadro, C. F. (1992). Levels of naturally occurring DNA polymorphism correlate with recombination rates in D. melanogaster. Nature, 356 (6369), 519–520. 10.1038/356519a0

Bonhomme, M., Boitard, S., San Clemente, H., Dumas, B., Young, N., & Jacquet, C. (2015). Genomic Signature of Selective Sweeps Illuminates Adaptation of Medicago truncatula to Root-Associated Microorganisms. Molecular Biology and Evolution, 32 (8), 2097–2110. 10.1093/molbev/msv092

Bourgeois, Y. X. C., & Warren, B. H. (2021). An overview of current population genomics methods for the analysis of whole-genome resequencing data in eukaryotes. Molecular Ecology, 30 (23), 6036–6071. 10.1111/mec.15989

Cao, M., Fu, Y., Guo, Y., & Pan, J. (2009). Chlamydomonas (Chlorophyceae) colony PCR. Protoplasma, 235 (1-4), 107–110. 10.1007/s00709-009-0036-9

Caron, D. A. (2009). Past President’s Address: Protistan Biogeography: Why All The Fuss? Journal of Eukaryotic Microbiology, 56 (2), 105–112. 10.1111/J.1550-7408.2008.00381.X

Casteleyn, G., Leliaert, F., Backeljau, T., Debeer, A.-E., Kotaki, Y., Rhodes, L., Lundholm, N., Sabbe, K., & Vyverman, W. (2010). Limits to gene flow in a cosmopolitan marine planktonic diatom. Proceedings of the National Academy of Sciences, 107 (29), 12952–12957. 10.1073/pnas.1001380107

Charron, G., Leducq, J.-B., & Landry, C. R. (2014). Chromosomal variation segregates within incipient species and correlates with reproductive isolation. Molecular Ecology, 23 (17), 4362–4372. 10.1111/mec.12864

Clark, R. M., Linton, E., Messing, J., & Doebley, J. F. (2004). Pattern of diversity in the genomic region near the maize domestication gene Tb1. Proceedings of the National Academy of Sciences of the United States of America, 101 (3), 700–707. 10.1073/pnas.2237049100

Craig, R. J., Böndel, K. B., Arakawa, K., Nakada, T., Ito, T., Bell, G., Colegrave, N., Keightley, P. D., & Ness, R. W. (2019). Patterns of population structure and complex haplotype sharing among field isolates of the green alga Chlamydomonas reinhardtii. Molecular Ecology, 28 (17), 3977–3993. 10.1111/MEC.15193

Craig, R. J., Gallaher, S. D., Shu, S., Salomé, P., Jenkins, J. W., Blaby-Haas, C. E., Purvine, S. O., O’Donnell, S., Barry, K., Grimwood, J., Strenkert, D., Kropat, J., Daum, C., Yoshinaga, Y., Goodstein, D. M., Vallon, O., Schmutz, J., & Merchant, S. S. (2022). The Chlamydomonas Genome Project, version 6: Reference assemblies for mating type plus and minus strains reveal extensive structural mutation in the laboratory. The Plant Cell, koac347. 10.1093/plcell/koac347

Culbertson, E. M., & Levin, T. C. (2023). Eukaryotic CD-NTase, STING, and viperin proteins evolved via domain shuffling, horizontal transfer, and ancient inheritance from prokaryotes. PLOS Biology, 21 (12), e3002436. 10.1371/journal.pbio.3002436

Custer, G. F., Bresciani, L., & Dini-Andreote, F. (2022). Ecological and Evolutionary Implications of Microbial Dispersal. Frontiers in Microbiology, 13. 10.3389/fmicb.2022.855859

Dale, R. K., Pedersen, B. S., & Quinlan, A. R. (2011). Pybedtools: A flexible Python library for manipulating genomic datasets and annotations. Bioinformatics, 27 (24), 3423–3424. 10.1093/BIOINFORMATICS/BTR539

De Wit, R., & Bouvier, T. (2006). “Everything is everywhere, but, the environment selects”; what did Baas Becking and Beijerinck really say? Environmental Microbiology, 8 (4), 755–758. 10.1111/j.1462-2920.2006.01017.x

Duan, G., Bao, J., Chen, X., Xie, J., Liu, Y., Chen, H., Zheng, H., Tang, W., & Wang, Z. (2021). Large-Scale Genome Scanning within Exonic Regions Revealed the Contributions of Selective Sweep Prone Genes to Host Divergence and Adaptation in Magnaporthe oryzae Species Complex. Microorganisms, 9 (3), 562. 10.3390/microorganisms9030562

Durante, L., Hübner, W., Lauersen, K. J., & Remacle, C. (2019). Characterization of the GPR1/FUN34/YaaH protein family in the green microalga Chlamydomonas suggests their role as intracellular membrane acetate channels. Plant Direct, 3 (6), e00148. 10.1002/pld3.148

Earl, D. A., & vonHoldt, B. M. (2012). STRUCTURE HARVESTER: A website and program for visualizing STRUCTURE output and implementing the Evanno method. Conservation Genetics Resources, 4 (2), 359–361. 10.1007/s12686-011-9548-7

Ellison, C. E., Hall, C., Kowbel, D., Welch, J., Brem, R. B., Glass, N. L., & Taylor, J. W. (2011). Population genomics and local adaptation in wild isolates of a model microbial eukaryote. Proceedings of the National Academy of Sciences, 108 (7), 2831–2836. 10.1073/pnas.1014971108

Evanno, G., Regnaut, S., & Goudet, J. (2005). Detecting the number of clusters of individuals using the software STRUCTURE: A simulation study. Molecular Ecology, 14 (8), 2611–2620. 10.1111/j.1365-294X.2005.02553.x

Excoffier, L., & Foll, M. (2011). Fastsimcoal: A continuous-time coalescent simulator of genomic diversity under arbitrarily complex evolutionary scenarios. Bioinformatics, 27 (9), 1332–1334. 10.1093/bioinformatics/btr124

Eydivandi, S., Roudbar, M. A., Karimi, M. O., & Sahana, G. (2021). Genomic scans for selective sweeps through haplotype homozygosity and allelic fixation in 14 indigenous sheep breeds from Middle East and South Asia. Scientific Reports, 11 (1), 2834. 10.1038/s41598-021-82625-2

Fenchel, T., & Finlay, B. J. (2003). Is microbial diversity fundamentally different from biodiversity of larger animals and plants? European Journal of Protistology, 39 (4), 486–490. 10.1078/0932-4739-00025

Fenchel, T., & Finlay, B. J. (2004). The Ubiquity of Small Species: Patterns of Local and Global Diversity. BioScience, 54 (8), 777–784.

Feng, C., Wang, J., Liston, A., & Kang, M. (2023). Recombination Variation Shapes Phylogeny and Introgression in Wild Diploid Strawberries. Molecular Biology and Evolution, 40 (3), msad049. 10.1093/molbev/msad049

Ferris, P. J., & Goodenough, U. W. (1997). Mating Type in Chlamydomonas Is Specified by mid, the Minus-Dominance Gene. Genetics, 146 (3), 859–869. 10.1093/genetics/146.3.859

Finlay, B. J., & Clarke, K. J. (1999). Apparent global ubiquity of species in the protist genus Paraphysomonas. Protist, 150 (4), 419–430. 10.1016/S1434-4610(99)70042-8

Flowers, J. M., Hazzouri, K. M., Pham, G. M., Rosas, U., Bahmani, T., Khraiwesh, B., Nelson, D. R., Jijakli, K., Abdrabu, R., Harris, E. H., Lefebvre, P. A., Hom, E. F. Y., Salehi-Ashtiani, K., & Purugganan, M. D. (2015). Whole-Genome Resequencing Reveals Extensive Natural Variation in the Model Green Alga Chlamydomonas reinhardtii. The Plant Cell, 27 (9), 2353–2369. 10.1105/tpc.15.00492

Foissner, W. (1999). Protist Diversity: Estimates of the Near-Imponderable. Protist, 150 (4), 363–368. 10.1016/S1434-4610(99)70037-4

Foissner, W. (2006). Biogeography and dispersal of micro-organisms: A review emphasizing protists. Acta Protozoologica, 45 (2), 111–136.

Foissner, W. (2008). Protist diversity and distribution: Some basic considerations. Biodiversity and Conservation, 17 (2), 235–242. 10.1007/S10531-007-9248-5/FIGURES/3

Ford, S. A., Craig, R. J., & Ness, R. W. (2022). A novel method for identifying Chlamydomonas reinhardtii (Chlorophyta) and closely related species from nature. Journal of Phycology. 10.1111/JPY.13306

Gallaher, S. D., Fitz-Gibbon, S. T., Glaesener, A. G., Pellegrini, M., & Merchant, S. S. (2015). Chlamydomonas Genome Resource for Laboratory Strains Reveals a Mosaic of Sequence Variation, Identifies True Strain Histories, and Enables Strain-Specific Studies. The Plant Cell, 27 (9), 2335–2352. 10.1105/tpc.15.00508

Gel, B., Díez-Villanueva, A., Serra, E., Buschbeck, M., Peinado, M. A., & Malinverni, R. (2016). regioneR: An R/Bioconductor package for the association analysis of genomic regions based on permutation tests. Bioinformatics, 32 (2), 289–291. 10.1093/bioinformatics/btv562

Gilbert, K. J. (2016). Identifying the number of population clusters with structure: Problems and solutions. Molecular Ecology Resources, 16 (3), 601–603. 10.1111/1755-0998.12521

Glenn, T. C., Nilsen, R. A., Kieran, T. J., Sanders, J. G., Bayona-Vásquez, N. J., Finger, J. W., Pierson, T. W., Bentley, K. E., Hoffberg, S. L., Louha, S., Garcia-De Leon, F. J., Del Rio Portilla, M. A., Reed, K. D., Anderson, J. L., Meece, J. K., Aggrey, S. E., Rekaya, R., Alabady, M., Belanger, M., … Faircloth, B. C. (2019). Adapterama I: Universal stubs and primers for 384 unique dual-indexed or 147,456 combinatorially-indexed Illumina libraries (iTru & iNext). PeerJ, 7, e7755. 10.7717/peerj.7755

Gorman, D. S., & Levine, R. P. (1965). Cytochrome f and plastocyanin: Their sequence in the photosynthetic electron transport chain of Chlamydomonas reinhardi. Proceedings of the National Academy of Sciencess of the United States of America, 54 (6), 1665–1669. 10.1073/pnas.54.6.1665

Hasan, A. R., Duggal, J. K., & Ness, R. W. (2019). Consequences of recombination for the evolution of the mating type locus in Chlamydomonas reinhardtii. New Phytologist, 224 (3), 1339–1348. 10.1111/nph.16003

Hasan, A. R., & Ness, R. W. (2020). Recombination Rate Variation and Infrequent Sex Influence Genetic Diversity in Chlamydomonas reinhardtii. Genome Biology and Evolution, 12 (4), 370–380. 10.1093/gbe/evaa057

Henden, L., Lee, S., Mueller, I., Barry, A., & Bahlo, M. (2018). Identity-by-descent analyses for measuring population dynamics and selection in recombining pathogens. PLOS Genetics, 14 (5), e1007279. 10.1371/journal.pgen.1007279

Hernandez, R. D. (2008). A flexible forward simulator for populations subject to selection and demography. Bioinformatics (Oxford, England), 24 (23), 2786–2787. 10.1093/bioinformatics/btn522

Hernandez, R. D., Kelley, J. L., Elyashiv, E., Melton, S. C., Auton, A., McVean, G., 1000 GENOMES PROJECT, Sella, G., & Przeworski, M. (2011). Classic Selective Sweeps Were Rare in Recent Human Evolution. Science, 331 (6019), 920–924. 10.1126/science.1198878

Hu, Y., Feng, C., Yang, L., Edger, P. P., & Kang, M. (2022). Genomic population structure and local adaptation of the wild strawberry Fragaria nilgerrensis. Horticulture Research, 9, uhab059. 10.1093/hr/uhab059

Huang, W., Guo, Y., Lysen, C., Wang, Y., Tang, K., Seabolt, M. H., Yang, F., Cebelinski, E., Gonzalez-Moreno, O., Hou, T., Chen, C., Chen, M., Wan, M., Li, N., Hlavsa, M. C., Roellig, D. M., Feng, Y., & Xiao, L. (2023). Multiple introductions and recombination events underlie the emergence of a hyper-transmissible Cryptosporidium hominis subtype in the USA. Cell Host & Microbe, 31 (1), 112–123.e4. 10.1016/j.chom.2022.11.013

Janes, J. K., Miller, J. M., Dupuis, J. R., Malenfant, R. M., Gorrell, J. C., Cullingham, C. I., & Andrew, R. L. (2017). The K = 2 conundrum. Molecular Ecology, 26 (14), 3594–3602. 10.1111/mec.14187

Ji, F., Ma, Q., Zhang, W., Liu, J., Feng, Y., Zhao, P., Song, X., Chen, J., Zhang, J., Wei, X., Zhou, Y., Chang, Y., Zhang, P., Huang, X., Qiu, J., & Pei, D. (2021). A genome variation map provides insights into the genetics of walnut adaptation and agronomic traits. Genome Biology, 22 (1), 300. 10.1186/s13059-021-02517-6

Ji, R., Edwards, M., Mackas, D. L., Runge, J. A., & Thomas, A. C. (2010). Marine plankton phenology and life history in a changing climate: Current research and future directions. Journal of Plankton Research, 32 (10), 1355–1368. 10.1093/plankt/fbq062

Kang, L., He, G., Sharp, A. K., Wang, X., Brown, A. M., Michalak, P., & Weger-Lucarelli, J. (2021). A selective sweep in the Spike gene has driven SARS-CoV-2 human adaptation. Cell, 184 (17), 4392–4400.e4. 10.1016/j.cell.2021.07.007

Kim, E.-K., Cho, D. H., Suh, S.-I., Lee, C.-J., Kim, H.-S., & Suh, H.-H. (2022). Isolation and Characterization of the Indigenous Microalgae Chlamydomonas reinhardtii K01 as a Potential Resource for Lipid Production and Genetic Modification. Journal of Life Science, 32 (3), 202–209.

Koc, A. (2021). Alkc/parallel-structure: Zenodo. 10.5281/zenodo.4697229

Korunes, K. L., & Samuk, K. (2021). Pixy: Unbiased estimation of nucleotide diversity and divergence in the presence of missing data. Molecular Ecology Resources, 21 (4), 1359–1368. 10.1111/1755-0998.13326

Kraemer, S. A., & Boynton, P. J. (2017). Evidence for microbial local adaptation in nature. Molecular Ecology, 26 (7), 1860–1876. 10.1111/mec.13958

Lawson, D. J., Hellenthal, G., Myers, S., & Falush, D. (2012). Inference of Population Structure using Dense Haplotype Data. PLOS Genetics, 8 (1), e1002453. 10.1371/journal.pgen.1002453

Leffler, E. M., Bullaughey, K., Matute, D. R., Meyer, W. K., Ségurel, L., Venkat, A., Andolfatto, P., & Przeworski, M. (2012). Revisiting an Old Riddle: What Determines Genetic Diversity Levels within Species? PLOS Biology, 10 (9), e1001388. 10.1371/journal.pbio.1001388

Li, H., & Durbin, R. (2009). Fast and accurate short read alignment with Burrows–Wheeler transform. Bioinformatics, 25 (14), 1754–1760. 10.1093/BIOINFORMATICS/BTP324

McKenna, A., Hanna, M., Banks, E., Sivachenko, A., Cibulskis, K., Kernytsky, A., Garimella, K., Altshuler, D., Gabriel, S., Daly, M., & DePristo, M. A. (2010). The Genome Analysis Toolkit: A MapReduce framework for analyzing next-generation DNA sequencing data. Genome Research, 20 (9), 1297–1303. 10.1101/GR.107524.110

McVean, G. (2009). A Genealogical Interpretation of Principal Components Analysis. PLOS Genetics, 5 (10), e1000686. 10.1371/journal.pgen.1000686

Merico, D., Isserlin, R., Stueker, O., Emili, A., & Bader, G. D. (2010). Enrichment Map: A Network-Based Method for Gene-Set Enrichment Visualization and Interpretation. PLOS ONE, 5 (11), e13984. 10.1371/journal.pone.0013984

Nakada, T., Shinkawa, H., Ito, T., & Tomita, M. (2010). Recharacterization of Chlamydomonas reinhardtii and its relatives with new isolates from Japan. Journal of Plant Research, 123 (1), 67–78. 10.1007/S10265-009-0266-0

O’Malley, M. A. (2008). “Everything is everywhere: But the environment selects”: Ubiquitous distribution and ecological determinism in microbial biogeography. Studies in History and Philosophy of Science Part C: Studies in History and Philosophy of Biological and Biomedical Sciences, 39 (3), 314–325. 10.1016/J.SHPSC.2008.06.005

Onetto, C. A., Sosnowski, M. R., Heuvel, S. V. D., & Borneman, A. R. (2022). Population genomics of the grapevine pathogen Eutypa lata reveals evidence for population expansion and intraspecific differences in secondary metabolite gene clusters. PLOS Genetics, 18 (4), e1010153. 10.1371/journal.pgen.1010153

Otasek, D., Morris, J. H., Bouças, J., Pico, A. R., & Demchak, B. (2019). Cytoscape Automation: Empowering workflow-based network analysis. Genome Biology, 20 (1), 185. 10.1186/s13059-019-1758-4

Pavlidis, P., & Alachiotis, N. (2017). A survey of methods and tools to detect recent and strong positive selection. Journal of Biological Research-Thessaloniki, 24 (1), 7. 10.1186/s40709-017-0064-0

Pritchard, J. K., Stephens, M., & Donnelly, P. (2000). Inference of population structure using multilocus genotype data. Genetics, 155 (2), 945–959. 10.1093/genetics/155.2.945

Puechmaille, S. J. (2016). The program structure does not reliably recover the correct population structure when sampling is uneven: Subsampling and new estimators alleviate the problem. Molecular Ecology Resources, 16 (3), 608–627. 10.1111/1755-0998.12512

Quinlan, A. R., & Hall, I. M. (2010). BEDTools: A flexible suite of utilities for comparing genomic features. Bioinformatics, 26 (6), 841–842. 10.1093/BIOINFORMATICS/BTQ033

Ramasamy, R. K., Ramasamy, S., Bindroo, B. B., & Naik, V. G. (2014). STRUCTURE PLOT: A program for drawing elegant STRUCTURE bar plots in user friendly interface. SpringerPlus, 3 (1), 431. 10.1186/2193-1801-3-431

Raudvere, U., Kolberg, L., Kuzmin, I., Arak, T., Adler, P., Peterson, H., & Vilo, J. (2019). G:Profiler: A web server for functional enrichment analysis and conversions of gene lists (2019 update). Nucleic Acids Research, 47 (W1), W191–W198. 10.1093/nar/gkz369

Rengefors, K., Kremp, A., Reusch, T. B. H., & Wood, A. M. (2017). Genetic diversity and evolution in eukaryotic phytoplankton: Revelations from population genetic studies. Journal of Plankton Research, 39 (2), 165–179. 10.1093/plankt/fbw098

Rohland, N., & Reich, D. (2012). Cost-effective, high-throughput DNA sequencing libraries for multiplexed target capture. Genome Research, 22 (5), 939–946. 10.1101/gr.128124.111

Sabeti, P. C., Reich, D. E., Higgins, J. M., Levine, H. Z. P., Richter, D. J., Schaffner, S. F., Gabriel, S. B., Platko, J. V., Patterson, N. J., McDonald, G. J., Ackerman, H. C., Campbell, S. J., Altshuler, D., Cooper, R., Kwiatkowski, D., Ward, R., & Lander, E. S. (2002). Detecting recent positive selection in the human genome from haplotype structure. Nature, 419 (6909), 832–837. 10.1038/nature01140

Salomé, P. A., & Merchant, S. S. (2019). A Series of Fortunate Events: Introducing Chlamydomonas as a Reference Organism. The Plant Cell, 31 (8), 1682–1707. 10.1105/TPC.18.00952

Santangelo, J. S., Johnson, M. T. J., & Ness, R. W. (2025). Signatures of selective sweeps in urban and rural white clover populations. Evolution, 79 (10), 2115–2132. 10.1093/evolut/qpaf138

Santangelo, J. S., Ness, R. W., Cohan, B., Fitzpatrick, C. R., Innes, S. G., Koch, S., Miles, L. S., Munim, S., Peres-Neto, P. R., Prashad, C., Tong, A. T., Aguirre, W. E., Akinwole, P. O., Alberti, M., Álvarez, J., Anderson, J. T., Anderson, J. J., Ando, Y., Andrew, N. R., … Johnson, M. T. J. (2022). Global urban environmental change drives adaptation in white clover. Science, 375 (6586), 1275–1281. 10.1126/science.abk0989

Schwartz, S. L., & Conn, G. L. (2019). RNA regulation of the antiviral protein 2*′*-5*′*-oligoadenylate synthetase. WIREs RNA, 10 (4), e1534. 10.1002/wrna.1534

Shannon, P., Markiel, A., Ozier, O., Baliga, N. S., Wang, J. T., Ramage, D., Amin, N., Schwikowski, B., & Ideker, T. (2003). Cytoscape: A Software Environment for Integrated Models of Biomolecular Interaction Networks. Genome Research, 13 (11), 2498–2504. 10.1101/gr.1239303

Škaloud, P., Jadrná, I., Dvořák, P., Škvorová, Z., Pusztai, M., Čertnerová, D., Bestová, H., & Rengefors, K. (2024). Rapid diversification of a free-living protist is driven by adaptation to climate and habitat. Current Biology, 34 (1), 92–105.e6. 10.1016/j.cub.2023.11.046

Spanner, R., Taliadoros, D., Richards, J., Rivera-Varas, V., Neubauer, J., Natwick, M., Hamilton, O., Vaghefi, N., Pethybridge, S., Secor, G. A., Friesen, T. L., Stukenbrock, E. H., & Bolton, M. D. (2021). Genome-Wide Association and Selective Sweep Studies Reveal the Complex Genetic Architecture of DMI Fungicide Resistance in Cercospora beticola. Genome Biology and Evolution, 13 (9), evab209. 10.1093/gbe/evab209

Stephan, W. (2019). Selective Sweeps. Genetics, 211 (1), 5–13. 10.1534/genetics.118.301319

Sundqvist, L., Keenan, K., Zackrisson, M., Prodöhl, P., & Kleinhans, D. (2016). Directional genetic differentiation and relative migration. Ecology and Evolution, 6 (11), 3461–3475. 10.1002/ece3.2096

Szpiech, Z. A. (2024). Selscan 2.0: Scanning for sweeps in unphased data. Bioinformatics, 40 (1), btae006. 10.1093/bioinformatics/btae006

Szpiech, Z. A., Novak, T. E., Bailey, N. P., & Stevison, L. S. (2021). Application of a novel haplotype-based scan for local adaptation to study high-altitude adaptation in rhesus macaques. Evolution Letters, 5 (4), 408–421. 10.1002/evl3.232

Voight, B. F., Kudaravalli, S., Wen, X., & Pritchard, J. K. (2006). A Map of Recent Positive Selection in the Human Genome. PLOS Biology, 4 (3), e72. 10.1371/journal.pbio.0040072

Weigand, H., & Leese, F. (2018). Detecting signatures of positive selection in non-model species using genomic data. Zoological Journal of the Linnean Society, 184 (2), 528–583. 10.1093/zoolinnean/zly007

Weir, B. S., & Cockerham, C. C. (1984). Estimating F-Statistics for the Analysis of Population Structure. Evolution, 38 (6), 1358–1370. 10.2307/2408641

Whittaker, K. A., & Rynearson, T. A. (2017). Evidence for environmental and ecological selection in a microbe with no geographic limits to gene flow. Proceedings of the National Academy of Sciences of the United States of America, 114 (10), 2651. 10.1073/pnas.1612346114

Zheng, X., Levine, D., Shen, J., Gogarten, S. M., Laurie, C., & Weir, B. S. (2012). A high-performance computing toolset for relatedness and principal component analysis of SNP data. Bioinformatics, 28 (24), 3326–3328. 10.1093/bioinformatics/bts606

